# Permissive epigenetic landscape facilitates distinct transcriptional signatures of activating transcription factor 6 in the liver

**DOI:** 10.1101/2021.03.10.434889

**Authors:** Anjana Ramdas Nair, Priyanka Lakhiani, Chi Zhang, Filippo Macchi, Kirsten C. Sadler

## Abstract

Proteostatic stress initiates a transcriptional response that is unique to the stress condition, yet the regulatory mechanisms underlying the distinct gene expression patterns observed in stressed cells remains unknown. Using a functional genomic approach, we investigated how activating transcription factor 6 (ATF6), a key transcription factor in the unfolded protein response (UPR), regulates target genes. We first designed a computational strategy to define Atf6 target genes based on the evolutionary conservation of predicted ATF6 binding in gene promoters, identifying 652 conserved putative Atf6 target (CPAT) genes. CPATs were overrepresented for genes functioning in the UPR, however, the majority functioned in cellular processes unrelated to proteostasis, including small molecule metabolism and development. Functional studies of stress-independent and toxicant based Atf6 activation in zebrafish livers showed that the pattern of CPAT expression in response to Atf6 overexpression, alcohol and arsenic was unique. Only 34 CPATs were differentially expressed in all conditions, indicating that Atf6 is sufficient to regulate a small subset of CPATs. Blocking Atf6 using Ceapins in zebrafish demonstrated that Atf6 is necessary for activation of these genes in response to arsenic. We investigated CPAT during physiologically mediated hepatocyte stress using liver regeneration in mice as a model. Over half of all CPATs were differentially expressed during this process. This was attributed to the permissive chromatin environment in quiescent livers on the promoters of these genes, characterized by the absence of H3K27me3 and enrichment of H3K4me3. Taken together, these data uncover a complex transcriptional response to Atf6 activation and implicate a permissive epigenome as a mechanism by which distinct transcriptional responses are regulated by Atf6.

## INTRODUCTION

Cells have an incredible capacity to orchestrate nuanced response to stimuli. This is best exemplified by the dynamic changes in gene expression that occur as cells traverse the initial response through adaptation and return to homeostasis following exposure to a toxicant. The maintenance of the cellular proteostasis environment is a well-studied process which integrates signaling pathways with changes in transcription and translation [1]. A wide range of environmental and physiological stimulants perturb the proteostasis environment, including stressors that directly impact the functioning of the endoplasmic reticulum (ER). Many stressors are pathological and the role of ER stress in many important diseases, including liver disease, cancer and neurodegenerative disorders, is extensively studied.

Sensing and countering this imbalance depends on a complex network of signaling pathways, transcription factors, chaperones and other factors collectively termed the unfolded protein response (UPR). Cells that fail to adequately respond to the unfolded protein burden in the ER can become dysfunctional; ER stress has been identified as a crucial molecular mechanism at the basis of a wide range of diseases, including liver disease [2–4]. UPR functions hinge on a transcriptional program which has traditionally been described as an upregulation of genes encoding proteins that function in the ER by the coordinated activity of three transcription factors: activating transcription factor 6 (ATF6), activating transcription factor 4 (ATF4) and X-box binding protein 1 (XBP1s) [5]. Despite the importance of these transcription factors, a well-defined target gene list has not been identified for any of them, and the mechanisms determining how these factors mount a nuanced response to different stressors is not known. This is, in part, due to the significant cross talk between the different UPR branches [6, 7] making it challenging to identify the transcriptional targets distinct to each branch.

The UPR becomes engaged in response to many disparate conditions, including pharmacological stressors, such as tunicamycin which blocks protein glycosylation, environmental toxicants which cause oxidative and ER stress, and pathological stressors, as occurs in response to environmental toxicants or a viral infection, as seen in the example of coronavirus infection which overwhelms the secretory pathway with the production of surface glycoproteins [8]. These responses induce a canonical UPR characterized by the upregulation of well-studied proteins such as the chaperones Bip (Hspa5) and Hyou1 or the ERAD factors Edem1 or Derl3. The majority of studies on UPR transcriptional targets has focused on only a few of these genes, with an emphasis on those that play a pivotal role in mitigating ER stress. Recent studies using transcriptomics to identify the patterns of gene expression accompanying ER stress have challenged this perception, revealing a complex UPR-associated transcriptomic pattern that does not follow a simple global transcriptional upregulation model [1, 9–12]. For instance, some studies have found downregulation of cell surface proteins and receptors as a consequence of UPR induction [10] while others showed that genes involved in mitochondrial function are downregulated in cells with ER stress [1]. Indeed, our work using zebrafish to study the relationship between ER stress and fatty liver has shown that genes are both up and down regulated in the context of UPR mediated fatty liver [13, 14]. However, such studies on the global transcriptional reprogramming in response to ER stress are underrepresented in the literature, and the regulation of non-canonical UPR mechanisms remain understudied. There is a gap in knowledge related to how the UPR transcriptome tailors its response to diverse stressors such as environmental toxicants, dietary intake, infection and in physiological processes such as development and regeneration.

ATF6 is the only UPR sensor that directly regulates the transcriptional response to ER stress. Unfolded proteins in the ER or presence of specific sphingolipids [15] facilitate ATF6 translocation to the Golgi apparatus where it is cleaved to release the transcriptional regulatory domain [16, 17]; this cleaved cytoplasmic domain (nATF6) translocates to the nucleus where it functions as a transcriptional regulator. However, neither the precise binding site nor the chromatin context required for the function of Atf6 as a transcriptional regulator have been well defined. One series of biochemical studies using HeLa cells identified the cis-regulatory elements, termed ER stress element (ERSE) I (CCAAT-(N)9-CCACG) and ERSE II (ATTGG-N-CCACG), as required for transcriptional activation of well characterized Atf6 target genes, such as Hspa5 [18–20]. These studies show that nAtf6 binds to CCACG sequence when the general transcription factor, NFY, is bound to the CCAAT site, and several genes that function in proteostasis have one of these sites in their proximal promoter [18, 20–23]. Another study identified TGACGTG as an Atf6 response element (Atf6 RE) to which human Atf6 binds *in vitro* [24]. A reporter construct with a modified version of the Atf6 RE in the context of a minimal promoter was used in a zebrafish model to show that this reporter gene is activated *in vivo* during ER stress in an Atf6 dependent manner [25]. This indicates that both the ERSE I and II sites as well as Atf6 RE can be used to identify Atf6 target genes. These conserved motifs have been found in the proximal promoters of several known ER stress response genes and UPR targets [18, 22, 23, 26]. *In vitro* studies have shown that Atf6 is both necessary and sufficient for the regulation of some of these canonical genes, including *HERPUD*, *DDIT3* (i.e. *CHOP*) and *XBP1*, [18, 20, 21, 26, 27] whereas the canonical target *HSPA5* (also called BiP) is regulated in an Atf6 dependent fashion in some settings, but not in others [28]. The recent study showing that Atf6 is activated by sphingolipids to drive metabolic genes whereas during proteostatic stress, Atf6 regulates ER chaperone genes [15]. How this switch in target selection by Atf6 is made is not clear.

Most transcription factors depend on distinct chromatin environments dictated by epigenetic modifications or cofactors to bind their consensus site. However, little is known about the impact of chromatin environment or the epigenetic regulation of any UPR genes. *In vitro* binding assays have revealed that NFY associates with nATF6, and the binding of NFY to ERSE facilitates recruitment of nATF6 to DNA. Other studies identified SRF [29] and YY1 [21, 30, 31] as ATF6 co-factors. In addition, a few studies have implicated epigenetic modifications in UPR target gene regulation [31–33]. However, it is not known if there are a set of genes for which Atf6 is necessary and sufficient for their regulation or if there are Atf6 target genes that require additional co-factors or specific chromatin environments. In fact, there is no clear consensus on what Atf6 target genes are.

While dysregulation of a single transcription factor may not be sufficient to modify the expression of all its target genes, it is necessary to identify during ER stress whether differential expression of putative ATF6 targets are due to ATF6 upregulation itself, or other factors such as either interaction with co-factors or changes in chromatin accessibility. The recent reports of drugs designed to modulate ATF6 activity [9, 28, 34–36] not only provide an exciting opportunity to treat diseases associated with disrupted proteostasis, but allows for comprehensive understanding of ATF6 targets and regulation. Our previous studies using zebrafish found that Atf6 was upregulated in response to various UPR activating toxicants such as tunicamycin, ethanol, and arsenic [13, 14, 37–41]. Despite some common features of the transcriptional response in these models, we noted distinct patterns of UPR-relevant gene expression in response to different stressors [14]. In addition, we found that transgenic overexpression of nAtf6 in zebrafish hepatocytes (*Tg(fabp10a:nAtf6-mCherry;cmlc2:EGFP)*) was sufficient to induce the expression of well-characterized Atf6 target genes involved in proteostasis in addition to genes involved in metabolism, which we found were, in part, involved in the fatty liver phenotype observed in this model [13]. Uncovering whether this phenotype is a direct result of Atf6 overexpression or is due to indirect consequences of stress-independent Atf6 modulation has important implications for understanding how Atf6 contributes to fatty liver disease. Interestingly, our previous work identified a cluster of ER-related genes such as *Derlin-3* and *Calreticulin* that are dynamically co-expressed during liver regeneration in response to partial hepatectomy (PH) in mice [42]. These genes, based on literature have been identified to be regulated in an Atf6-dependent manner. This inspired us to investigate the role of Atf6 in the physiological demanding process of liver regeneration.

In this study, we use computational approaches to identify a list of Atf6 targets based on evolutionary conservation of putative Atf6 binding sites in gene promoters. We verified these targets *in vivo* using zebrafish to define how Atf6 target gene expression is modulated in the liver. Surprisingly, few of these genes were classified as playing a role in the UPR. Instead, most are classified as functioning in developmental processes and metabolism, indicating that Atf6 has roles in addition to regulating proteostasis. RNAseq data from an *in vivo* system where nAtf6 is overexpressed identified a subset of these targets whereby Atf6 is sufficient for their regulation; these include canonical UPR genes widely used to assess Atf6 activation, confirming these as *bona fide* UPR targets. By examining the regulation of target genes when Atf6 is activated by disease relevant environmental exposures (ethanol, arsenic) and detailed analysis of the shared and distinct profiles of target gene activation across these biological contexts we identified a small set of Atf6-sufficient genes were differentially expressed in response to each of these stressors. However, the pattern of expression of these genes was unique to each condition, reflecting distinct Atf6 mediated transcriptomic signatures in response to different challenges. To examine the response of the liver to cell stress mediated by a physiological challenge, we examined the expression of Atf6 targets during liver regeneration in mice. We found that over half of Atf6 targets were dynamically expressed during this process and discovered a specific chromatin landscape in quiescent livers was the distinguishing feature between Atf6 targets that were differentially expressed and the ones that did not change during regeneration. These data uncover a role for epigenetic modifications in dictating which targets are selected by this important transcription factor.

## MATERIAL AND METHODS

### Atf6 Target Gene Identification

The proximal promoter (500 bp upstream of the transcription start site (TSS)) was acquired for all transcripts in *Danio rerio, Homo sapiens*, and *Mus musculus* from BioMart using the ‘upstream_flank’ filter for all Ensembl transcript IDs. These promoters were filtered for each species to identify those that did not overlap with other annotated elements such as coding regions or promoters of other genes. These unique promoters were filtered for containing exact match of previously reported ATF6 binding sites by searching for 3 different sequences: 1. the first half of the ERSE I site (NFY binding site, CCAAT) or the ERSE II site, which is its reverse complement (ATTGG), 2. the Atf6 consensus sequence (CCACG) or 3. the Atf6 response element (Atf6 RE; TGACGTG) in each species. The unified set of all these elements contained 652 genes that contained either an ERSEI or ERSEII site or the Atf6 RE in their proximal promoters were termed as CPATs. To validate this dataset we used publicly available datasets that have been functionally verified and published from HEK293 cells where Atf6 was overexpressed using an inducible system (i.e. genes where Atf6 expression is sufficient for regulation) [4] and from the liver of Atf6^-/-^ mice treated with tunicamycin to identify those genes that failed to be differentially expressed due to their requirement for Atf6 [43]. We identified significantly differentially expressed genes in either condition and asked whether CPATs were included in these datasets.

### UPR Gene Identification

A unified list of genes involved in the Unfolded Protein Response was assembled by combining human genes annotated with the UPR Reactome Pathway, Response to ER Stress (GO:0034976), and Response to Unfolded Protein (GO:0006986). Human GO and Reactome annotations were used since functional annotation of zebrafish genes is considerably less extensive; GO annotations were filtered by Experimental Evidence codes to exclude genes annotated without manual curation. 37 genes with more than one ortholog in the human genome were removed resulting in the final list of 314 UPR genes in zebrafish.

### Animal Maintenance and Treatment

Adult fish were maintained at a 14:10 light:dark cycle at 28°C. The transgenic line *Tg(fabp10:nAtf6-mCherry-C; cmlc2:GFP)-C* was previously described [13]. Fertilized embryos from group matings were reared in plates of 60 with embryo water (0.6 g/L Crystal Sea MarineMix; Marine Enterprises International, Baltimore, MD) containing Methylene Blue at 28°C. *Tg(fabp10:nAtf6-mCherry-C; cmlc2:GFP)-C* transgenic fish were screened for transgenesis based on GFP expression in the heart at 3 days post fertilization (dpf), and non-transgenic siblings were used as a control. In the case of ethanol treatment, embryos were exposed to 30 ml solutions of 2% ethanol in embryo water from 96-120 hours post fertilization (hpf). Plates were sealed with Parafilm to prevent evaporation. In the case of arsenic treatment, embryos were exposed to 10 ml solutions of 1.5mM iAs in embryo water in 6-well plates from 96-120 hours post fertilization (hpf). All procedures were performed in accordance with the New York University Abu Dhabi Institutional Animal Care and Use Committee (IACUC).

Ceapins A7 (a gift from the Walter Lab) was used to block Atf6 activation in zebrafish larvae. We treated 15 zebrafish larvae at 80 hpf with 4.0 uM Ceapins or DMSO as a control. At 96 hpf, samples were segregated into 4 treatment groups so that DMSO or Ceapins treated larvae were exposed to 1.0 mM or were left unchallenged until 120 hpf when they were assessed for morphological abnormalities and survival. The larvae were then anaesthetized with 500 μM tricaine (Ethyl 3-aminobenzoate methanesulfonate; Sigma-Aldrich) and livers were microdissected for RNA extraction.

### Image Acquisition

For whole mount imaging of live larvae, embryos were anesthetized with 500 μM tricaine (Ethyl 3-aminobenzoate methanesulfonate; Sigma-Aldrich), mounted in 3% methyl-cellulose on a glass slide and imaged on a Nikon SMZ25 stereomicroscope.

### Quantitative real time PCR (qPCR)

Livers were microdissected from 5 dpf zebrafish larvae anesthetized using tricane and immobilized in 3% methyl cellulose. 10-12 livers were collected per sample and processed for RNA extraction using TRIzol Reagent as previously described [44]. RNA was reverse transcribed using the qScript cDNA SuperMix, and SYBR Green Fast Mix. Samples were analyzed in triplicate on QuantStudio 5 (Thermo Fisher), with at least 2 independent samples for each experiment. Target gene expression was normalized to *rpp0* expression and expressed relative to either the DMSO or Ceapin controls from the same clutch via the comparative threshold cycle method. Primer sequences are listed in Table S1.

### RNAseq

Total RNA was extracted from 15-20 livers dissected from zebrafish larvae for each sample at 78 and 120 hpf. For nAtf6 overexpression dataset, 2 clutches at 78 hpf and 3 clutches at 120 hpf of *Tg(fabp10:nAtf6-mCherry-C)* larvae were used for RNA extraction (GSE130800 and GSE130801). For arsenic treatment, 5 clutches of larvae were exposed to 1.5 mM iAs from 96–120 hpf as described ([41] and GSE156419). For EtOH treatment, larvae were exposed to 350 mM ethanol from 96–120 hpf (GSE151291). cDNA libraries were sequenced on the Illumina HiSeq2500 platform to obtain 100 bp paired-end reads. Sequencing quality was assessed by using FASTQC [45] and the reads were quality trimmed using Trimmomatic [46] to remove low quality reads and adapters. Reads were aligned to the *Danio rerio* GRCz10 reference genome assembly with HISAT2 [47]. To estimate gene expression, read counts were calculated by HTSeq [48] with the ‘union’ mode and ensemble annotation [49]. A generalized linear model, which was implemented in DESeq2 in Bioconductor [50], was adopted to test differential gene expression. Adjusted p-value (FDR) <0.05 was treated as significantly different expression. RNAseq datasets from mouse samples collected from 7 time points from male mouse livers after PH (GSE125006) were previously described [42].

### ATAC-seq

100 mg of liver tissue was homogenized with a tissue Dounce and 50,000 nuclei were used to prepare libraries as previously described (48) checked for correct size distribution with bioanalyzer. Sequencing was performed on Hiseq2500 platform for 100 cycles single end read runs at the Genomics Core facility of the NYUAD. Sequenced reads were aligned to the mouse reference genome (GRCm38.p4) with BWA-MEM, generating sorted and cleaned bam with PICARD for further analysis.

### ATAC-seq and ChIPseq analysis

We utilized previously published ChIPseq datasets for H3K4me3 and H3K27me3 obtained from the liver of 6-8 week old male mice (GSE125006) that were previously described [42]. We identified regions around TSS of mouse CPATs that were differentially expressed at any time point in the RNAseq datasets obtained following PH and compared them to the CPATs that were unchanged. We visualized the ATACseq peak intensity (+/- 2kb around TSS), Chipseq peak intensity of H3K4me3 and H3K27me3 (+/-3kb around TSS) using EnrichedHeatmap package in R/bioconductor package and generate the metaplotsmeta [51]. To compare gene expression of CPATs that change versus those that remain unchanged in mouse livers were assessed based on a p-Value cut off for >0.05 (CPATs that did not change) or p<0.05 (CPATs that changed). Gene expression was normalizednormalized as log (FPKM+1).

### Statistical Analysis

Normalization and test of differential gene expression in RNAseq datasets were conducted in R using the ‘DESeq2’ package using a generalized linear model. GO enrichment analysis was conducted using the GO hypergeometric over-representation test in the ‘ClusterProfiler’ package in R using default parameters, and REVIGO was subsequently used to eliminate redundant enriched terms. The bias in directionality of differentially expressed CPATs was assessed using the non-parametric Mann-Whitney U test. Survival plots and qPCR data were plotted using Graphpad Prism.

## RESULTS

### Identification of conserved putative Atf6 target genes

To identify all the possible target genes of nAtf6, we reasoned that genes that are regulated by Atf6 would contain one of the previously defined Atf6 binding sites (ERSE I or II or the Atf6 RE) in their proximal promoters and that this binding site would be conserved across vertebrates. To validate this approach, we scanned 200 bp upstream of transcription start site (TSS) of zebrafish, mouse and human homologs of two known Atf6 target genes *hspa5* [19, 52, 53] and *herpud1* [26] for the presence of canonical binding sites. Both the Atf6 RE and ERSE I was found in the *hspa5* promoter and an ERSE I was found in the *herpud1* promoter, albeit with some divergence between zebrafish and mammalian sequences (Figure 1A). We expanded this to develop a genome wide strategy to identify all Conserved Putative Atf6 Target genes (CPATs) using inclusion criteria in which genes have (1) an exact match of the ERSE I, ERSE II or Atf6 RE within 500bp of their proximal promoters and (2) orthologous genes in human, mouse and zebrafish genes (Figure 1B). Genes where the proximal promoters overlapped with another genomic element, *i.e.* the 3’UTR or the promoter of another gene, were excluded. Non-overlapping promoters were selected and screened for an exact match of one of the following sequences: (1) the NFY binding site CCAAT (or it’s reverse complement ATTGG) which is one feature of ERSE I and II, (2) the Atf6 binding site in ERSE I and ERSE II (CCACG) or (3) the Atf6 RE (TGACGTG) to generate 3 different lists of putative target genes for each species. We then identified the homologous genes present in all 3 datasets for each list and retained the zebrafish homolog as the reference (Table S2). This approach identified 5,259 zebrafish genes with the NFY site, 1,011 with the ERSE I/II Atf6 consensus site and 17 with the Atf6 RE (Figure 1B and Table S2). There were 637 genes that had both components of the ERSE I or ERSE II sites, 2 of which also had the Atf6 RE (*cad* and *pax6a*), and 15 additional genes with only the Atf6 RE. The unified set of conserved genes containing ERSE I or II and Atf6 RE in the promoters was identified as 652 zebrafish CPATs (Figure 1B and Table S3).

**Figure 1.**
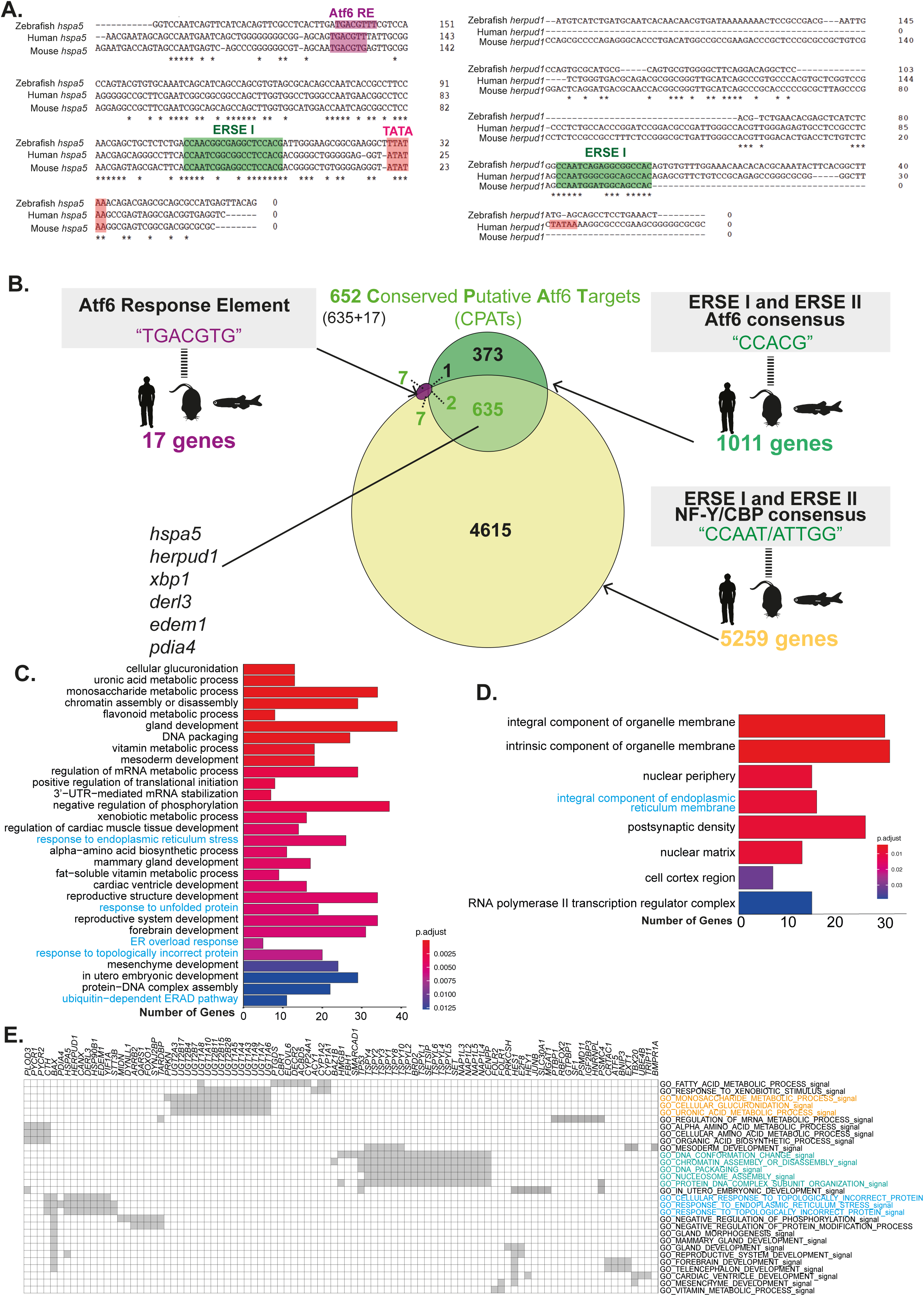
Curation and functional annotation of CPATs. (A) Sequence alignment of promoter regions (200bp upstream of TSS) of *hspa5* and *herpud1* across human, mouse and zebrafish. Highlighted regions indicate ERSE I in green, Atf6 RE in pink and TATA in red. (B) Filtering strategy to identify CPATs included 3 criteria: (i) the first half of the ERSE I and II consensus sequence required for NFY binding: CCAAT/ATTGG (ii) the second half of the ERSE I and II shown to have Atf6 binding activity: CCACG and (iii) the Atf6 RE shown to have Atf6 binding activity: TGACGTG. Sequence pertaining to each criteria was identified in the proximal promoters (acquired from BioMart, designated as −500 bp from the TSS) of orthologous genes in zebrafish, mouse, and human. Venn diagram indicates overlap between each criteria. Cumulative set of genes either having a complete ERSEI/II site : 635 genes, the Atf6 RE: 15 genes or both: 2 were designated as constitutive putative Atf6 target genes or CPATS : 652 genes. *Bona fide* CPATs are shown in black (C) Gene Ontology (GO) based biological process enrichment analysis of top 25 terms in putative Atf6 targets; terms related to proteostasis highlighted in blue (D) GO cellular component analysis : terms related to ER highlighted in blue. (E) Geneset enrichment Leading Edge analysis highlights the groups of genes driving a particular function. Grey boxes mark the genes representing each GO term. Highlighted in blue are proteostasis, orange are small molecule metabolism and green are DNA packing related GO terms.

We compared this strategy to two alternative computational approaches to predict Atf6 target genes. We used the R package ‘tftargets’ and the software Harmonize (http://amp.pharm.mssm.edu/Harmonizome/gene_set/ATF6/TRANSFAC+Curated+Transcription+Factor+Targets) to identify potential Atf6 targets. ‘tftargets’ identified 1,022 targets and Harmonize identified a geneset of 116 genes curated by functional studies. Importantly, neither of these datasets included the canonical Atf6 target genes, such as *hspa5*, *herpud1*, *hyou1* or *derl3*. Moreover, gene ontology (GO) enrichment analysis for putative targets generated from tftargets included terms related to lipid metabolic process, Golgi vesicle transport and vesicle organization, but not to ER stress or proteostasis (Figures S1B). Due to minimal overlap between the two gene lists and the low representation of known Atf6-related biological processes, we excluded these datasets for the following analysis.

To validate CPATs by functional studies, we used well characterized genetic models in which Atf6 was modulated and for which there are publicly available microarray datasets: (1) human embryonic kidney (HEK293T) cells engineered to have inducible overexpression of nAtf6 [9] and (2) the liver of *Atf6a* knock out mice assessed 8 and 24 hours following ER stress induced by exposure to tunicamycin [35]. The overexpression of nAtf6 identified 41 genes categorized as uniquely regulated by Atf6. In contrast, Atf6a deficiency in mice did not affect the expression of any genes in the untreated liver. However, 5,216 genes were differentially expressed compared to wild-type (WT) mice at either 8 or 24 hours after tunicamycin injection (Figure S2A and Table S4). We found a significant overlap between CPATs and the genes identified in mouse, but not with the human cells. This could reflect a lack of conservation of the consensus sites in these genes in other species or could indicate the inclusion of genes that are indirect targets of Atf6.

To determine whether CPATs were grouped into functional categories, we used the Gene Ontology (GO) Overrepresentation Test with the human CPAT homologs since the human genome has a better functional annotation compared to zebrafish. As expected, GO biological function terms related to proteostasis and the response to ER stress were enriched, but surprisingly, the majority of the terms encompassed genes involved in small molecule metabolism, including xenobiotics, vitamins and amino acids, and those related to development, differentiation and morphogenesis (Figure 1C). Analysis of the cellular component of CPATs leads to a similar conclusion: while the ER membrane was highly enriched, the cell cortex and nuclear structures were also included (Figure 1D). Moreover, GO categories related to the response to ER stress and the UPR were enriched in CPATs and in the human and mouse genesets that identified putative Atf6 targets based on functional studies (Figures S2A-B). Additionally, several of the metabolic process enriched in CPATs overlapped with the GO terms enriched in the dataset from Atf6 knock-out mice [43] (Figure S2C). This suggests that a conserved function of Atf6 is to regulate genes that are involved in metabolism and development.

We used Gene Set Enrichment Leading Edge Analysis to assess whether our analysis was influenced by genes common to several GO terms and thus artificially expanded the GO terms associated with these genesets. We found 3 clusters of genes that were linked to multiple GO terms, one that was dominated by UDP glucuronosyltransferases, the second was dominated by genes that encode nuclear proteins, such as the testes specific protein Y encoded (TSYs) and nucleosome assembly proteins (NAPs), and a third by genes involved in the UPR (Figure 1E). It is possible that inclusion of multiple members of a gene family arises from the conversion between the zebrafish to human orthologs, nevertheless these analyses suggest that several GO terms that are dominated by CPAT genes are redundant. In contrast, GO terms related to developmental processes are populated by unique set of genes, indicating these are not redundant. This is in line with studies that have uncovered a role for Atf6 in development and tissue homeostasis [25, 54, 55]. Taken together, the curated list of CPATs extend the role of Atf6 in cellular processes beyond proteostasis, namely xenobiotic metabolism and nuclear organization.

### A Subset of CPATs Are Involved in the UPR

Atf6 is best studied for its role as a transcriptional regulator of genes that function in the UPR and this was confirmed by the GO analysis of CPATs (Figure 1C-E). We investigated this further by assessing whether CPATs were overrepresented in a geneset encompassing all genes that function in the UPR. We identified and combined all human genes annotated with GO terms ‘Response to ER Stress’ and ‘Response to Unfolded Protein,’ and the Reactome pathway ‘Unfolded Protein Response’ to generate a unified dataset (Table S5). The zebrafish homologs of these genes were then identified, discarding those with more than one paralogous zebrafish gene (Table S5) to generate a zebrafish UPR geneset containing 314 genes (Figure 2A). Of these, 23 overlapped with CPATS, including the *hspa5*, *herpud1*, *edem1* and *derl3* which are genes that have been functionally verified as Atf6 targets in other systems (Figure 2B; Table S6) [18, 20, 21, 26, 27]. These data indicate that a very small proportion (4.2%) of CPATs are annotated as playing a role in the UPR, and that only 8.9% of all UPR genes contained a conserved Atf6 binding site. This is consistent with data in Figure 1C indicating that Atf6 may regulate genes that participate in other cellular functions that extend beyond the UPR. This also confirms previous studies indicating that multiple transcription factors regulate UPR genes [9, 56–58] which could account for the low overlap between UPR genes and CPATs.

**Figure 2.**
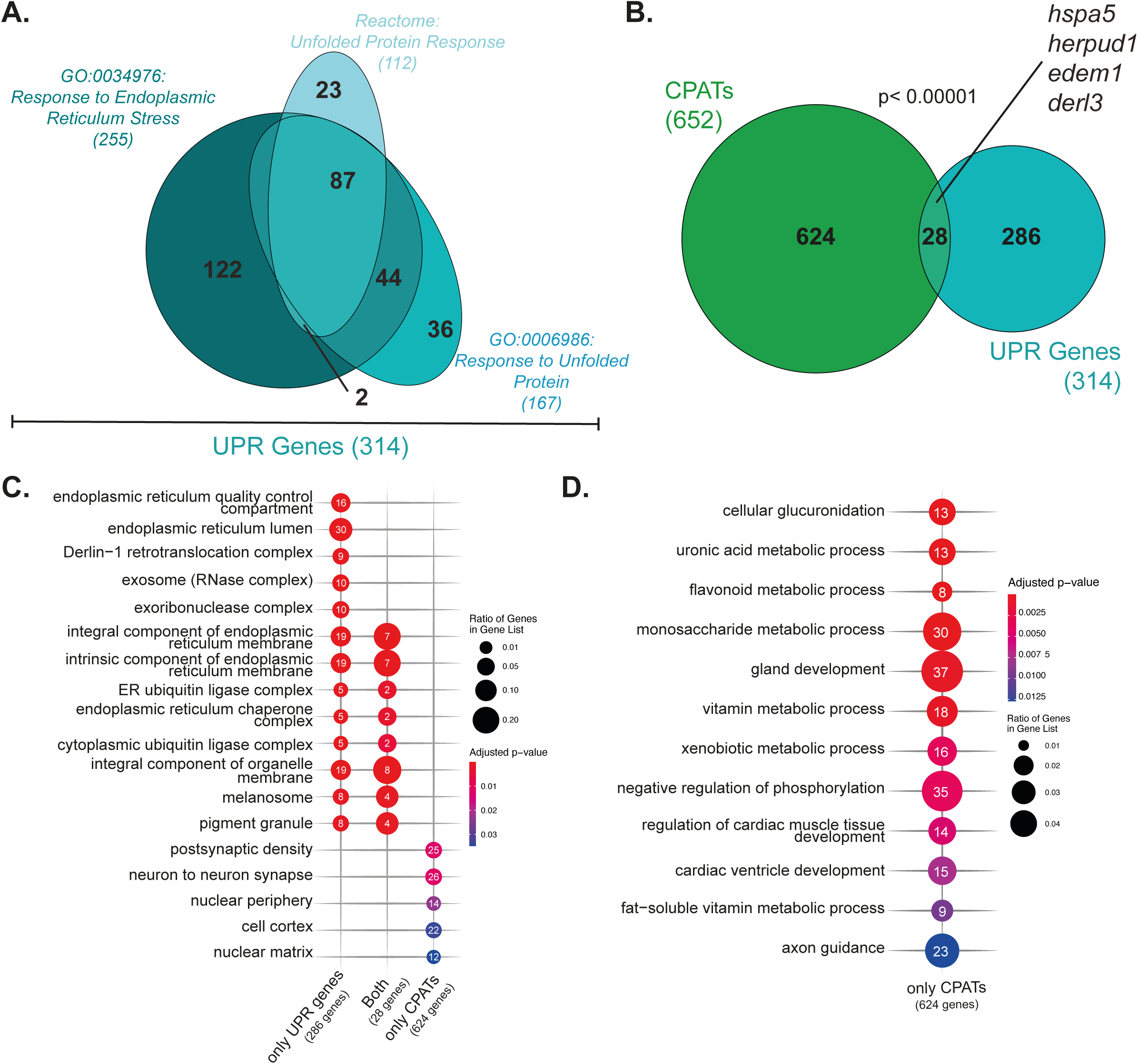
A minority of CPATs are involved in proteostasis. (A) Venn diagram depicting overlap of UPR Genes annotated to REACT_18356, GO:0034976, and GO:0006986 in humans obtained from Reactome and AmiGO respectively. (B) Human UPR genes were converted to zebrafish homologs (turquoise) for integration with the CPATs (green) with *bona fide* CPATs shown in black. Significant enrichment of CPATs in UPR gene dataset was determined by Fisher’s exact test. (C) & (D) Gene ontology-based cellular component or biological process overrepresentation analysis of overlapping and non-overlapping CPATs and UPR genes and of CPATs that are not UPR targets; terms related to proteostasis are blue. Statistically significant results are plotted with point size depicting fraction of genes in each gene list annotated to each GO term; point numbers showing the exact number of genes annotated.

We reasoned that genes with functions in the UPR would be localized to the ER while those CPATs with functions distinct from proteostasis may be localized to other cellular compartments. GO classification of cellular compartments showed that, as expected, UPR genes largely localized in the ER lumen and in complexes that are formed in the ER, whereas the CPATs that were not UPR genes localized elsewhere, suggesting that Atf6 regulates also genes that function outside the ER (Figure 2C). GO enrichment of the biological processes of the non-UPR CPATs revealed these roles outside the ER are related to development and metabolism (Figure 2D). This is consistent with functional studies implicating Atf6 in development of bone, cartilage, eye, the nervous system and other tissues [54] and uncovers potentially novel roles for Atf6 regulating energy and small molecule metabolism.

### nAtf6 Overexpression is Sufficient to Regulate a Subset of CPATs

To validate those CPATs that could be regulated by Atf6 directly, we used an *in vivo* functional approach. Since UPR is relevant for liver pathology and Atf6 activation has an important role in fatty liver, we focused our studies on the liver [13, 59]. We used a transgenic zebrafish line where exons 2-10 of the Atf6 gene encoding the nAtf6 domain (Figure S3A) is overexpressed in hepatocytes (*Tg(fabp10a:nAtf6-mCherry;cmlc2: GFP*) [13]. RNAseq analysis of two independently-generated RNAseq datasets (GSE130801, Figure S3B-C) of the nAtf6 overexpressing livers show increased transcriptional levels of the exons encoding the nAtf6 domain compared to their wild-type siblings at 120 hpf. Both qPCR (Figure S3D) and standard reverse transcriptase-PCR (Figure S3E) using primers that detect the transgene sequence as well as the endogenous *atf6* transcript (Fq1 and Rq1 in Figure S3A) show a significant increase in the amount of *atf6* transcript in the liver of these transgenics. As a control, we confirmed that elevated levels of the transgene were not attributed to DNA contamination (Figure S3E). In addition, we also show that expression of the well-characterized downstream target gene, *hspa5*, was significantly increased in these samples, confirming this CPAT as a direct Atf6 target (Figure S3D). We previously showed that the nAtf6-mCherry fusion protein was expressed in adult livers using Western blotting [13]. Here, we extend this analysis in 120 hpf livers of *Tg(fabp10:nAtf6-mCherry; cmlc2:GFP)-C* larvae and demonstrate the presence of the fusion protein, albeit at low levels (Supplementary Figure S3F) possibly due to the endogenous mechanisms that serve to degrade the Atf6 protein [60, 61]. These data validate and extend our published report, indicating 2-5 fold increase of nAtf6 expression over endogenous levels in this transgenic line [13], indicating that independent and skilled investigators generate highly reproducible results from this model and establishes this as useful tool to examine the effects of modest nAtf6 overexpression on target gene expression.

To identify genes directly regulated by nAtf6 in stress-independent conditions, we analyzed RNAseq from *Tg(fabp10:nAtf6-mCherry; cmlc2:GFP)-C* livers and their non-transgenic siblings at 78 hpf (2 samples) and 120 hpf (3 samples) (Figure 3A). The early developmental time point was chosen as it is estimated to be within 40 hours from the first expression of the *fabp10a* driven transgene [62] and the earliest time in which reliable liver dissection could be performed. The second time point (120 hpf) was used to identify genes that require prolonged nAtf6 interaction for their regulation. In the 78 hpf dataset, there were 184 significantly differentially expressed genes (DEGs) based on a p value <0.05, with 17 downregulated and 167 upregulated (Figure S4 A-B). There was significant overlap of these DEGs with the putative Atf6 target genes identified through functional studies in mouse samples (Figure S4C). At 120 hpf, there were 4,614 significantly differentially expressed genes (DEGs) based on a p value <0.05, with 1,844 downregulated and 2,770 upregulated (Figure S4 D-E). Moreover, there was significant overlap of the DEGs with those that were differentially expressed when Atf6 was manipulated in mouse and human cells (Figure S4 C,F). By 120 hpf when the liver is mature, nearly all CPATs were expressed (FPKM > 5) in the liver of wild-type siblings (Figure S4G), consistent with the fact that hepatocytes are a highly secretory cell type and that Atf6 is expressed in the liver of WT larvae (Figure S3B-C). Almost all DEGs observed at 78 hpf were differentially expressed at 120 hpf (Figure 3B). This indicates that Atf6 overexpression sustains the regulation of its target genes for over 36 hours *in vivo*.

**Figure 3.**
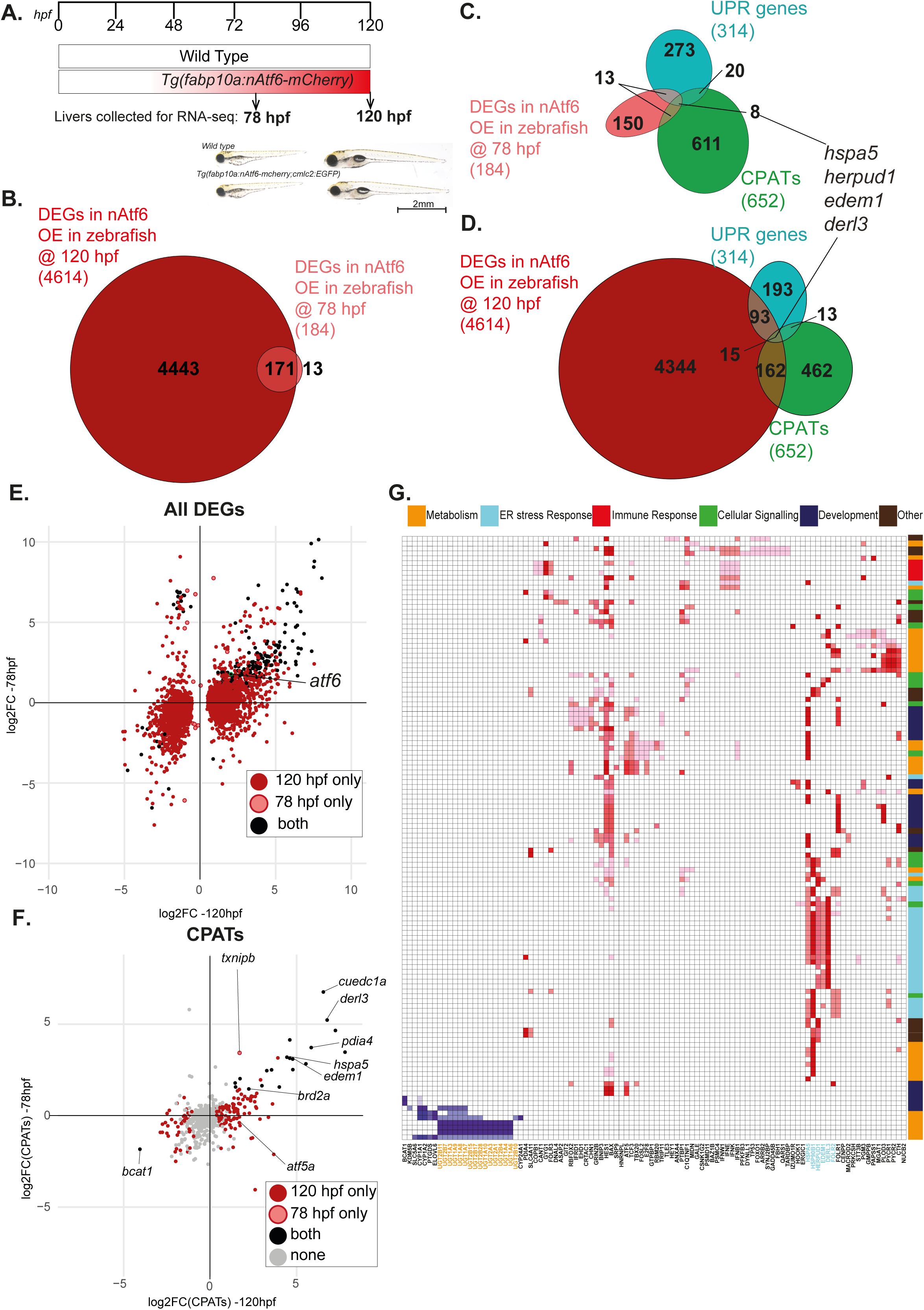
Stress-independent nAtf6 overexpression activates a small subset of putative Atf6 or UPR target genes. (A) Experimental set up of RNA-seq liver collections at 78 hpf and 120 hpf wild type zebrafish larvae and *Tg(fabp10a:nAtf6-mcherry;cmlc2:GFP)-C* transgenics used in these studies. Representative images of wild type zebrafish larvae and *Tg(fabp10a:nAtf6-mcherry;cmlc2:GFP)-C* transgenics at 78hpf and 120hpf. (B) Venn diagram depicting the overlap between genes significantly differentially expressed (p<0.05) in nAtf6 over expressing zebrafish livers at 78hpf (pink) and 120hpf (dark red). (C-D) Venn diagram depicting overlap of CPATs(green), UPR genes (turquoise), and DEGS in nAtf6 at (C) 78 hpf (pink) and (D) 120hpf (dark red). (E) Cross plot showing the DEGS unique to 120 hpf (dark red), unique to 78hpf (pink with a red circle) and those that are differentially expressed in both (black). Atf6 indicated by black circle with green circle. (F) Cross plot showing CPATs unique to 120 hpf (dark red), unique to 78hpf (pink with a red circle) and those that are differentially expressed in both (black). Grey indicates CPATs that do not change. (G) GSEA Leading edge analysis showing GO terms pertaining to sufficient CPATs that are upregulated and downregulated during nAtf6 over expression.

If Atf6 is sufficient to regulate the expression of CPATs, then CPATs should be overrepresented in the RNAseq datasets from the liver of *Tg(fabp10:nAtf6-mCherry; cmlc2:GFP)-C* larvae. Among the 184 DEGs detected in the 78 hpf dataset, 11.4% (21 genes) were CPATs, which we refer to as immediate response CPATs (IR-CPATs) (Figure 3C). In addition, 21 DEGs in these samples were UPR genes (Figure 3B), 8 of which were included among IR-CPATs. These consisted of known Atf6 targets that function in UPR such as *hspa5*, *herpud1*, *edem1*, *derl3* and *pdia4*. At 120 hpf, 177 of 4,614 DEGs were CPATs (Table S7), representing the 3.8% of all DEGs at this time point (177/4614) and 27% of all CPATs (177/652; Figure 3D; Table S7). Among these 177 CPATs, 20 of them were IR-CPATs. Hence forth, we referred to the 157 CPATs that were uniquely differentially expressed at 120 hpf as delayed response CPATs (DR-CPATs). Since these 177 CPATs were differentially expressed upon Atf6 overexpression under stress-free conditions, we termed these *Sufficient* CPATs, meaning that the presence of nAtf6 alone is sufficient to change their expression in the liver. Nearly 35% of all UPR genes (108/314) were differentially expressed in response to nAtf6 overexpression at 120 hpf, with 15 of the UPR genes classified as *Sufficient* CPATs (Figure 3D). These findings show that the majority of UPR genes are not differentially regulated in the liver in response to stress-independent activation of Atf6 and only a fraction of the *Sufficient* CPATs are involved in the UPR. This suggests that other transcriptional regulators are required to regulate UPR genes in stress free conditions.

To test if the stringent criteria for CPAT identification based on evolutionary conservation excluded genes which are regulated in by Atf6 in zebrafish, we investigated the proximal promoters of the zebrafish genes alone for the presence of ERSE I, II or Atf6 RE. This analysis uncovered additional 2 DEGs at 78 hpf and 304 DEGs at 120 hpf with Atf6 putative binding sites (Table S7). These genes were not included initially among the CPATs due to the lack of orthologs in either mouse or human, however this does not dramatically enhance the representation of CPATs among the DEGs in stress free Atf6 overexpression conditions in zebrafish.

Almost 70% (475/652) CPATs were not differentially expressed in this system and we defined them as *Insufficient CPATs* (Figure 3B-D; Table S7). This indicates that the majority of CPATs are either not accessible for regulation in the liver due to their chromatin environment or that other factors contribute to the regulation of CPATs in addition to Atf6. We reasoned that insufficient CPATs were either false positives from our screening method or that regulation of these genes by Atf6 requires presence of cofactors or changes in chromatin accessibility which are not present in the conditions examined here.

The UPR functions in large part by inducing the expression of chaperones and other genes involved in protein folding [1,3,4]. This suggests that the Atf6 and the other transcription factors regulating this response act as transcriptional activators. We found a trend towards upregulation (log_2_ Fold Change > 1, p value< 0.05) of DEGs at both 78 hpf and 120 hpf, but we noted that there were also genes which were downregulated in each dataset in *Tg(fabp10:nAtf6-mCherry; cmlc2:GFP)-C* samples (Figure 3E; S4A-B,D-E and Table S7). We further confirmed that the trend for upregulation was maintained among the *sufficient* CPATs, with majority of the canonical CPATs that function in UPR such as *hspa5*, *derl3*, *pdia4* and *edem1* were significantly upregulated at both time points (Figure 3F). Among the 45 CPATs that are downregulated in nAtf6 overexpressing samples at 120 hpf, only 1 is a UPR gene. This confirms that nAtf6 acts predominantly as a transcriptional activator for genes involved in proteostasis. However, our data strongly suggest that Atf6, like other bZip proteins, may act as a transcriptional repressor on some targets or could indirectly modulate the chromatin landscape to downregulate the expression of CPATs.

### Atf6 Overexpression induces Transcriptional Regulators and Cellular Processes other than Proteostasis

Our data showed that nAtf6 overexpression induces a significant group of genes that were not CPATs (4443 DEG at 120 hpf +13 additional DEG at 78 hpf −177 sufficient CPATs=4,279). We defined them as *Indirect Atf6 targets*, meaning that they change their expression as a consequence of the cellular environment induced by Atf6 overexpression (Figure 3B-D; Table S7). We hypothesized that *Indirect Atf6 targets* could be secondary targets regulated by CPATs. To test this, we first identified the human genes that corresponded to the *Indirect Atf6 targets* and used these to perform Ingenuity Pathway Analysis (IPA). This identified the top canonical pathways affected by stress-free overexpression of nAtf6 in zebrafish livers as the UPR, immune response, signaling, metabolism and other pathways that appear unrelated to proteostasis, with ATF4, XBP1, TNF, EGFR as hubs in this network (Figure S5A). This suggests that the *Indirect Atf6 targets* could be regulated by these transcriptional networks.

We next asked what other biological processes were impacted by sufficient CPATs and *Indirect Atf6 targets*. The most significantly-enriched GO terms for sufficient CPATs and indirect targets were similar; they were mainly involved in proteostasis and small molecule metabolism (Figure S5B). We noted that the clustering of GO terms representing closely related processes (i.e. uronic acid, glycoronate and xenobiotic metabolism) were driven by the redundancy of these GO terms due to a shared set of genes across all: the UDP glucuronyltransferases are significantly downregulated in these samples as indicated by geneset enrichment leading edge analysis (Figure 3E). Moreover, the GO terms involved in proteostasis are similarly highlighted by a core cluster of genes (Figure 3E). This suggests that Atf6 mediates a pattern of downstream target genes that have distinct functional roles and they are implicated in diverse biological pathways, as shown by others [15]. In summary, this analysis revealed that nAtf6 overexpression in zebrafish livers in stress-independent conditions is sufficient to differentially regulate over 1/3 of CPATs, suggesting the presence of other mechanisms, such co-factors or chromatin accessibility, that could contribute to CPAT regulation even in the presence of Atf6.

### The Cellular Response to Environmental Exposures Shapes the Atf6-mediated Transcriptional Response

To explore how the cellular environment influences the ability of Atf6 to regulate CPATs, we examined CPAT expression during environmental exposures that induce hepatocyte stress in the liver. To address this, we generated RNAseq datasets of the livers of zebrafish exposed to 350 mM ethanol (EtOH; 4 clutches; Figure 4A) or 1 mM inorganic arsenic (iAs; 5 clutches; Figure 4B). These were chosen because they are clinically relevant as significant contributors to liver disease worldwide and they have been shown to induce the expression of *atf6* and canonical Atf6 targets in the liver [13, 14, 38, 40, 41, 63, 64]. Both samples have significant *atf6* and *hspa5* induction in the liver (Figure S5A and Table S7), verifying Atf6 activation in these models.

**Figure 4.**
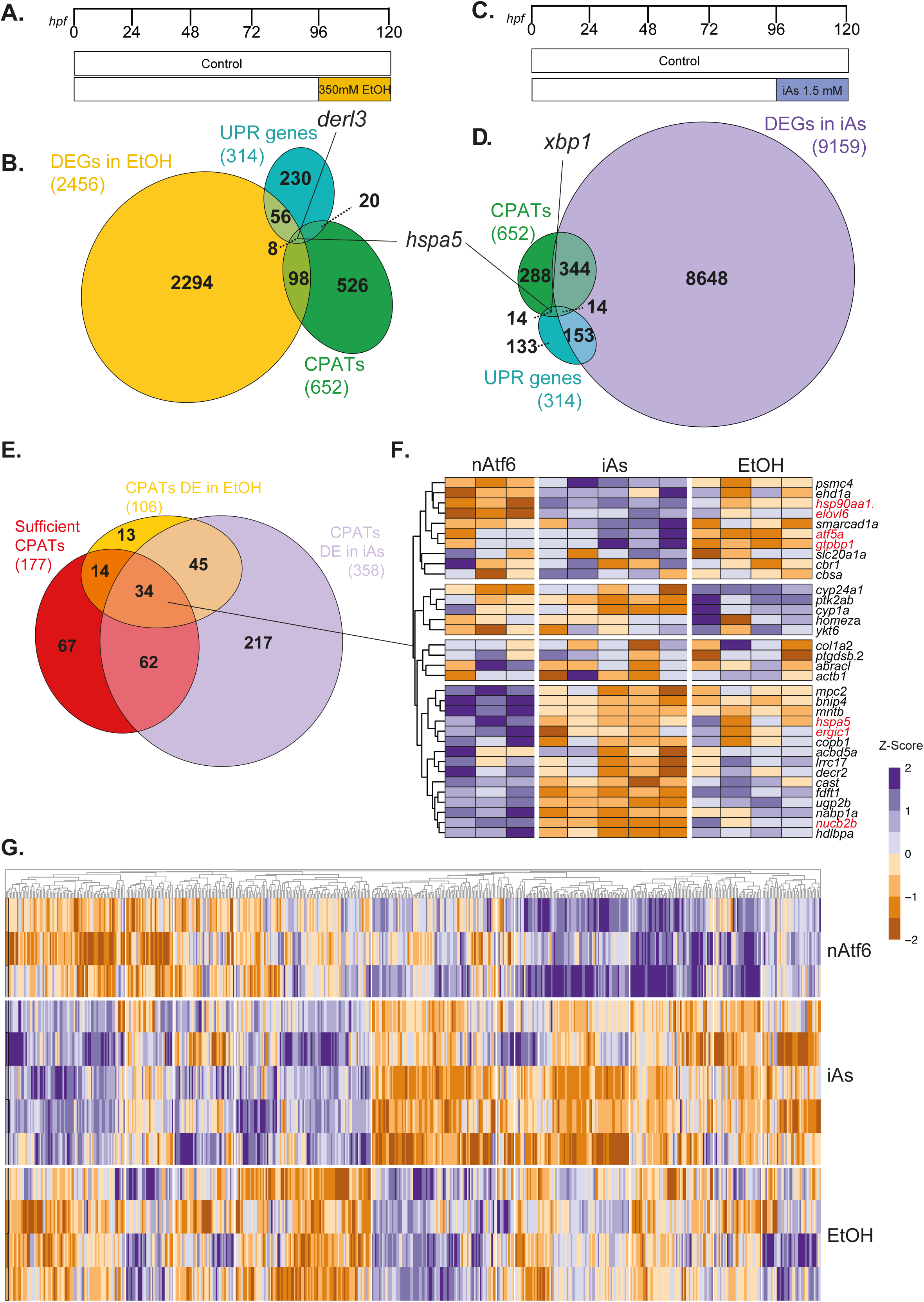
Stress-free nAtf6 expression and Toxicant-induced stress regulate distinct transcriptional signatures in the liver. A) EtOH exposure scheme showing zebrafish larvae exposed from 96-120 hpf with 350 mM EtOH and the livers were dissected and processed for RNAseq at 120 hpf (B) Venn diagram depicting the overlap between significantly DEGs in EtOH conditions (yellow), CPATs (green) and UPR genes (turquoise) (C) iAs exposure scheme showing zebrafish larvae exposed to 1 mM iAs from 96-120 hpf. (D) Venn diagram depicting the overlap between significantly DEGs in iAs conditions (violet), Atf6 targets (green) and UPR genes (turquoise). (E) Venn diagram depicting the overlap of CPATs differentially expressed in EtOH (yellow), iAs (violet) and sufficient CPATs defined in figure 3. (red). (F&G) Heatmap showing unsupervised clustering of the 34 common CPATs (F) and all CPATs(G) in nAtf6, iAs and EtOH conditions based on z-score.

EtOH exposure resulted in 2,456 DEGs (1,195 up and 1,261 down; p value <0.05; Figure 4C; Table S7. Less than 5% of DEGs (106 genes) were CPATs, representing 16% of all CPATs, a lower number than found in stress independent Atf6 overexpression conditions (Figure 3B). Of all DEGs, 64 were categorized as functioning in the UPR, representing 2.6% of all DEGs and 20% of all UPR genes; 8 of the UPR genes were also CPATs (Figure 4B). This shows that EtOH induces the UPR, and confirms our previous finding that ER stress is one of the cellular responses to alcoholic liver disease [13, 38, 64, 65].

Exposure to iAs resulted in 9,159 DEGs (4,135 up and 5,024 down; p value<0.05; Figure 4D; Table S7); among which were 358 CPATs that represent 54% of all CPATs and 167 UPR genes, (53% of all UPR genes; Figure 4D). Of these UPR genes, 14 were also CPATs (Figure 4D). This extends our previous finding that iAs activates the UPR in the liver [41, 63].

We used these 3 datasets – nAtf6 overexpression, EtOH and iAs treated – to identify those CPATs differentially expressed in each condition separately and those that differentially expressed in all conditions. There was a complex pattern of unique and shared differentially expressed CPATs in these datasets (Figure 4E), which demonstrates the nuanced transcriptional response to ER stress. For instance, among the CPATs that were differentially expressed in stress free conditions of Atf6 overexpression (i.e. sufficient CPATs), 48 were also differentially expressed response to EtOH and 96 were differentially expressed in response to iAs; of these, only 34 CPATs were common to all 3 datasets (Figure 4E). A similar analysis of the UPR genes showed that only 19 were shared between all datasets (Figure S7A), reiterating that there are distinct subclasses of the UPR depending on the stressor [14]. This suggests a differential Atf6 response in this setting of iAs and EtOH induced cell stress response.

GO enrichment analysis was performed to identify the biological processes impacted by stress independent nAtf6 expression and stress conditions, EtOH and iAs. The 34 CPATs common to all 3 datasets were involved in small molecule metabolism, and the CPATs that were not changed in any condition have a role in development and chromatin organization. Interestingly, the CPATs that were changed in stress-free nAtf6 overexpression samples were involved in proteostasis, protein transport and signaling, whereas the CPATs that were uniquely changed in the iAs samples functioned in process taking pace in the nucleus. This further confirms that the role of Atf6 extends beyond the ER stress response, and it is interesting to speculate that these genes may be regulated by Atf6 in other biological conditions, such as in distinct cell types or developmental contexts.

We were particularly interested in the 34 CPATs that were common to all 3 datasets and whether they had the same transcriptional response across all three conditions, i.e did they follow the same expression trend regardless of how nAtf6 was induced. To our surprise, unsupervised clustering revealed there was unique expression pattern of these CPATs in each condition, and not a single gene showed the same expression pattern in all 3 datasets. For examples, one cluster of genes that included *hspa5* were induced in by nAtf6 overexpression were suppressed in response to iAs and showed modest induction by EtOH. Another cluster of genes which included the metabolic genes *elovl6*, *cbr1* and *cbsa*, were suppressed in stress free nAtf6 overexpression and EtOH exposure, but strongly induced by iAs. Unsupervised clustering of the expression pattern of all CPATs confirmed this pattern was also consistent across all CPATs (Figure 4G). In summary, while nAtf6, EtOH and iAs exposure activate and drive expression of distinct CPATs in the liver, there is a clear differential response, indicating that the stimuli induced specific cellular environment may influence the Atf6-mediated transcriptional output. Taken together, these results indicate that Atf6 tailors its response depending on the stress condition, that it can function as both a transcriptional activator or repressor, and that only a small set of genes are always regulated when Atf6 is present. This reinstates previous work from our lab, that UPR responses are tailored dependent on stress condition [14].

### The Inhibition of Atf6 Activation Blocks CPAT Induction in Response to iAs

To determine if the differential expression of CPATs in response to iAs required Atf6, we used Ceapins A7 to block Atf6 activation [66]. Zebrafish larvae were pretreated from 80-96 hpf with 4.0 µM Ceapins or DMSO as a control, and then were subdivided into a control arm with no additional treatment or challenged with 1.0 mM iAs as a co-treatment from 96-120 hpf (Figure 5A). While Ceapins alone had no discernable effects on survival or overt phenotypes, it sensitized larvae to iAs treatment based on increased incidence of larvae with morphological defects such as spinal curvature, edema, irregular heartbeat, distended gut and brain necrosis (darkening of the head) and decreased overall survival (Figure 5B-C). This highlights the protective role of Atf6 during iAs stress response and that lack of Atf6 mediated transcriptional response leads to worse outcome in iAs-treated fish. We selected three groups of CPATs to examine by qPCR: those classified as Atf6 targets based on the literature (*hspa5, pdia4, der1*), those included in the 34 common CPATs (i.e differentially expressed in all 3 conditions examined in Figure 4), or those that were upregulated only in response to iAs exposure (iAs unique CPATs). We then selected genes that were upregulated in response to iAs exposure as these would be the most amenable to analysis by qPCR (Table S8). We evaluated the expression of a subset of genes in the liver of larvae in each condition (controls, Ceapins alone, iAs alone and Ceapins + iAs). We also assessed Atf6 expression, which, as expected, was not impacted by Ceapins, since this compound acts to inhibit Atf6 protein [67]. qPCR analysis confirmed that all the genes selected were significantly induced by iAs (Table S8; Figure 5D) and Ceapins treatment alone had no significant effect on gene expression (Table S8). However, 5 out of the 6 common CPATs, and 2 of the 4 iAs-unique CPATs were significantly suppressed by Ceapin treatment, demonstrating that these genes require Atf6 for their induction in response to iAs (Figure 5D and Table S8). In contrast, none of the canonical Atf6 targets were suppressed by Ceapins, suggesting that compensatory mechanisms may control these key UPR genes in response to iAs. This shows that Atf6 is required for the differential expression of the majority of CPATs that are induced in the liver by iAs exposure.

**Figure 5.**
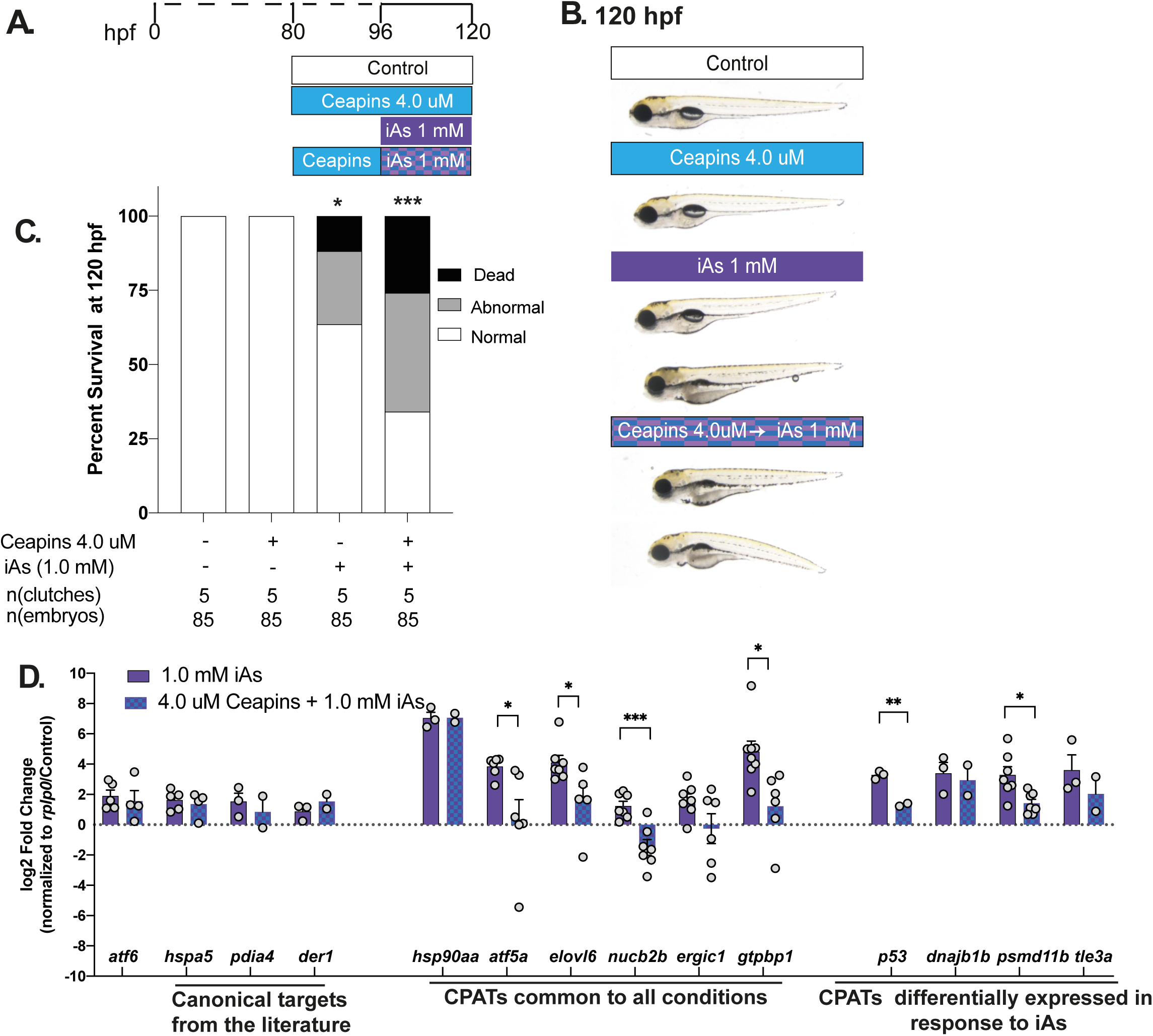
Atf6 inhibition by Ceapins A7 suppresses CPAT expression during acute iAs exposure. (A) Exposure scheme showing zebrafish larvae were pre-treated with 4.0 uM Ceapins from 80hpf and then co-exposed to 1mM iAs and Ceapins from 96-120hpf. (B) Representative images of 120hpf zebrafish larvae subjected to treatments in (A). Arrow highlight the phenotypes such as spinal curvature, edema, distended gut and brain necrosis (darkening of the head) compared to controls. (C) Percent of normal, abnormal and dead zebrafish at 120hpf per treatment condition; n=85, 5 clutches; *** p<0.0005, *p<0.05, by two-way ANOVA. (D) Gene expression analysis by qPCR depicting log2 Fold change of select genes categorized as CPATs based on literature, commonly DE in all three conditions and uniquely DE in iAs. (**** p<0.00005, **p<0.005, *p<0.05, by unpaired t-test)

### CPATs are differentially expressed during liver regeneration in mice

Nearly all of the previously published work on Atf6 focuses on its role in response to proteotoxic stress caused by exposure to a toxic substance, as in our studies on iAs and EtOH. We were interested in determining if Atf6 functions during liver regeneration based on our previous work where we identified a cluster of ER-related genes are differentially expressed in response to partial hepatectomy (PH) in mice [42]. Since these genes included *sufficient* CPATs, we used this as a model to examine CPATs expression in a physiological challenge that is unrelated to toxicant exposure.

We analyzed a series of RNAseq datasets generated from wild-type (WT) male mice quiescent livers and from 6 time points following PH (24, 30, 40, 48, 96 and 168 hours)[42]. These times were chosen to reflect distinct stages of this process in which synchronous reentry of hepatocytes into the cell cycle restores liver mass by 5 days after PH [68]. We found that *Atf6* levels increased between 24-48 hours after PH, and downregulated at 96 hours and 7 days after PH (Figure 6A). To assess CPATs in this process, we converted zebrafish genes to the mouse homologs, resulting in 744 mouse CPAT genes. Over half (52%; 397/744) of mouse CPATs were significantly differentially expressed at one or more time points following PH (Figure 6B and Table S9), far exceeding the extent of changes in response to toxicant exposure or Atf6 overexpression in zebrafish (Figure 5A) or in mice (Figure S2B). Among these 397 genes, 52 genes were sufficient CPATs (Table S10).

**Figure 6.**
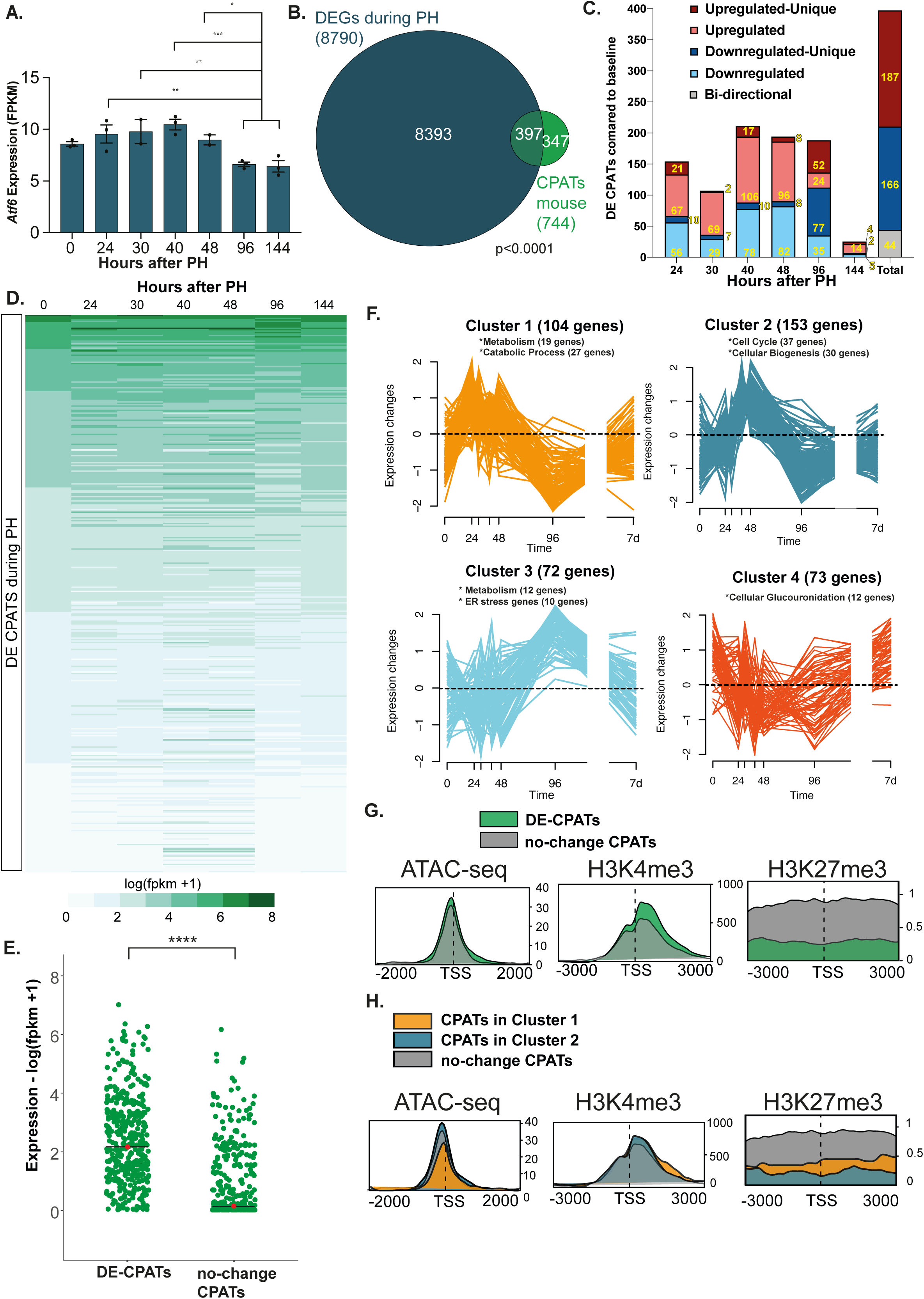
H3K27me3 promoter occupancy differentiates CPAT expression during mouse liver regeneration. (A) *Atf6* expression based on RNAseq at different timepoints in regenerating mouse liver after partial hepatectomy. (*** p<0.0005, **p<0.005, *p<0.05, by two-tailed t-test). (B) Venn diagram depicting overlap between DEGs during PH and CPATs converted to mouse genes. p<0.0001 based on Chi-square test. (C) Bar graph showing distribution of CPAT expression in regenerating control livers. Yellow numbers indicate the number genes in each subset. (D) Heatmap showing expression based on log(fpkm +1) of DE CPATs sorted based on baseline expression in regenerating control livers. (E) Jitter plot comparing expression between DE-CPATs and no-change-CPATs in quiescent mouse livers. **** indicates p<0.00005 by Mann-Whitney Test. (F) Gene clusters of co-regulated genes identified by unsupervised clustering of DE CPATs in regenerating control livers. The total number of gens in each cluster and those in the top GO categories are noted. (G) Averaged line profiles of ATAC-seq and Chip-seq of H3K4me3 and H3K27me3 between DE-CPATs (green) and no-change CPATs (grey). (H) Averaged line profiles of ATAC-seq and Chip-seq of H3K4me3 and H3K27me3 between CPATs in Cluster 1(orange) and 2(blue) and that do not change (grey).

CPATs showed a dynamic pattern of expression changes after PH, where some are changing their expression at only one time point (i.e. unique genes in Figure 6C), others at multiple time points, while 44 genes are found to be upregulated at some time points and downregulated at others (bidirectional; Figure 6C and Table S10). Most notably, a significant proportion of CPATs are downregulated at 96 hours after PH, corresponding to the time when Atf6 is also downregulated (Figure 6A). Genes that were expressed at the highest levels remain highly expressed during regeneration, while those that were expressed at lower levels increased expression (Figure 6D). Moreover, those genes that were dynamically expressed during regeneration were expressed at significantly higher levels than those that did not (Figure 6D-E). We reasoned that CPATs that were regulated by Atf6 presence would follow a similar pattern of expression during regeneration and therefore could be used to identify co-regulated genes. Unsupervised clustering of CPATs that were differentially expressed during regeneration showed that the majority of CPATs (402) were associated with distinct expression patterns (Figure 6F and Table S10). Clusters 1 & 2 are characterized by genes that peak in expression between 30-48 hours and then are downregulated at 96 hours: Cluster 1 is overrepresented for genes involved in metabolism and Cluster 2 is overrepresented for genes that function in the cell cycle. Cluster 3 genes show an opposite pattern, with expression peaking at 96 hours and are overrepresented for genes involved in metabolism and ER stress (Figure 6F). Interestingly, the CPATs that are silenced in the quiescent liver are enriched for genes involved in chromatin organization and neuronal processes (Figure S8). These data indicate that Atf6 targets sharing similar biological functions are co-expressed during regeneration, suggesting a common mechanism of regulation.

### CPAT expression during regeneration reflects the chromatin environment

Epigenetic modifications are a central mechanism for regulating transcription factor access. We hypothesized that the differential CPATs expression during regeneration could be influenced by the presence of activating or repressive epigenetic marks. To test this, we examined previously generated datasets mapping the epigenetic landscape in quiescent livers [42]. These included trimethylated histone H3 lysine 4 (H3K4me3) which marks active genes, H3K27me3 which is a repressive mark and ATACseq, which provides information about chromatin accessibility. We asked whether the chromatin environment was related to the expression pattern of CPATs in quiescent livers and during regeneration. We found that the majority of the CPATs that did not change following PH (‘no change’ CPATs) were completely silenced at baseline (244 of 347 no-change CPATs had FPKM < 5; Figure 6E). We expected these to be in a repressed chromatin environment compared to those that were differentially expressed (DE-CPATs). Surprisingly, there was no striking difference in accessibility surrounding the transcription start site (TSS) of no-change CPATs compared to DE-CPATs (Figure 6F) but we did see a marked enrichment and a broader peak of H3K4me3 on DE-CPATs compared to no-change CPATs. Interestingly, H3K27me3 was enriched on the TSS of no-change CPATs compared to DE-CPATs, suggesting this repressive mark as a mechanism for restraining Atf6 access to these genes during regeneration. We predicted that CPATs in cluster 1 and 2 would be the most accessible to Atf6 regulation since they are induced at early time points during regeneration. This was validated by the finding that, compared to no-change CPAT, cluster 1 and 2 CPATs were enriched for H3K4me3 downstream of the TSS and H3K27me3 was absent (Figure 6G). These data suggest that the ability of Atf6 to regulate target genes in the liver is dictated by the epigenetic environment. This creates a model whereby CPATs genes are accessible to Atf6, but some are suppressed by H3K27me3. It is interesting to note that silenced CPATs were also occupied by H3K4me3, suggesting these may be poised for activation in response to a different condition in which Atf6 is activated. Together, these data suggest that the chromatin environment provides important cues that dictate the differential CPATs expression signatures in response to different stressors.

## DISCUSSION

This study addresses how a highly nuanced transcriptional response can be orchestrated in the liver in response to diverse cellular challenges. The response to ER stress in vertebrates integrates multiple signaling pathways that converge on a transcriptomic response which we [14] and others [9, 35] have shown to be context dependent. Little is known about how the central UPR regulator, Atf6, selects target genes, or further, what those targets are. We provide the first systems level analysis of the Atf6 transcriptional response *in vivo*. The strategy used is a novel and comprehensive, albeit highly stringent, approach to identify putative Atf6 target genes. Similar bioinformatic approaches have been used successfully for other transcription factors or DNA-binding proteins to identify downstream motifs or genes of interest [69, 70]. We focused on the liver because there is a clear functional role for UPR regulators in liver biology and pathology [3, 4, 12, 13, 40, 43, 71, 72] and our initial presumption was that Atf6 would primarily function to regulate the response to ER stress. We were surprised that only a subset of the 652 CPATs we identified were annotated as playing a role in proteostasis. This is supported by other studies [15, 54, 73–75] and by our analysis of the gene lists generated from other approaches to identify Atf6 targets, which revealed that the majority of putative targets function in diverse cellular processes. We were also surprised that a minority of CPATs were differentially expressed both in datasets from other models where Atf6 expression was manipulated and in the datasets we generated in zebrafish from stress independent and toxicant exposure conditions [13, 14, 40, 44, 63, 64]. Together, these data show an Atf6-directed transcriptomic signature specific for each condition, with a small subset of CPATs with coincident expression patterns across the three models. We verified that the majority of these genes required Atf6 activity by showing that Ceapins A7 treatment effectively suppressed expression of iAs-induced CPATs. We identified the novel role of Atf6 and its targets during mouse liver regeneration and demonstrated the impact of repressive marks in CPAT regulation. Taken together, these data validate this integrated functional genomic approach to identify new CPAT genes and implicate Atf6 in biological processes that have not previously been investigated such as small molecule metabolism and development (Figure 7).

**Figure 7.**
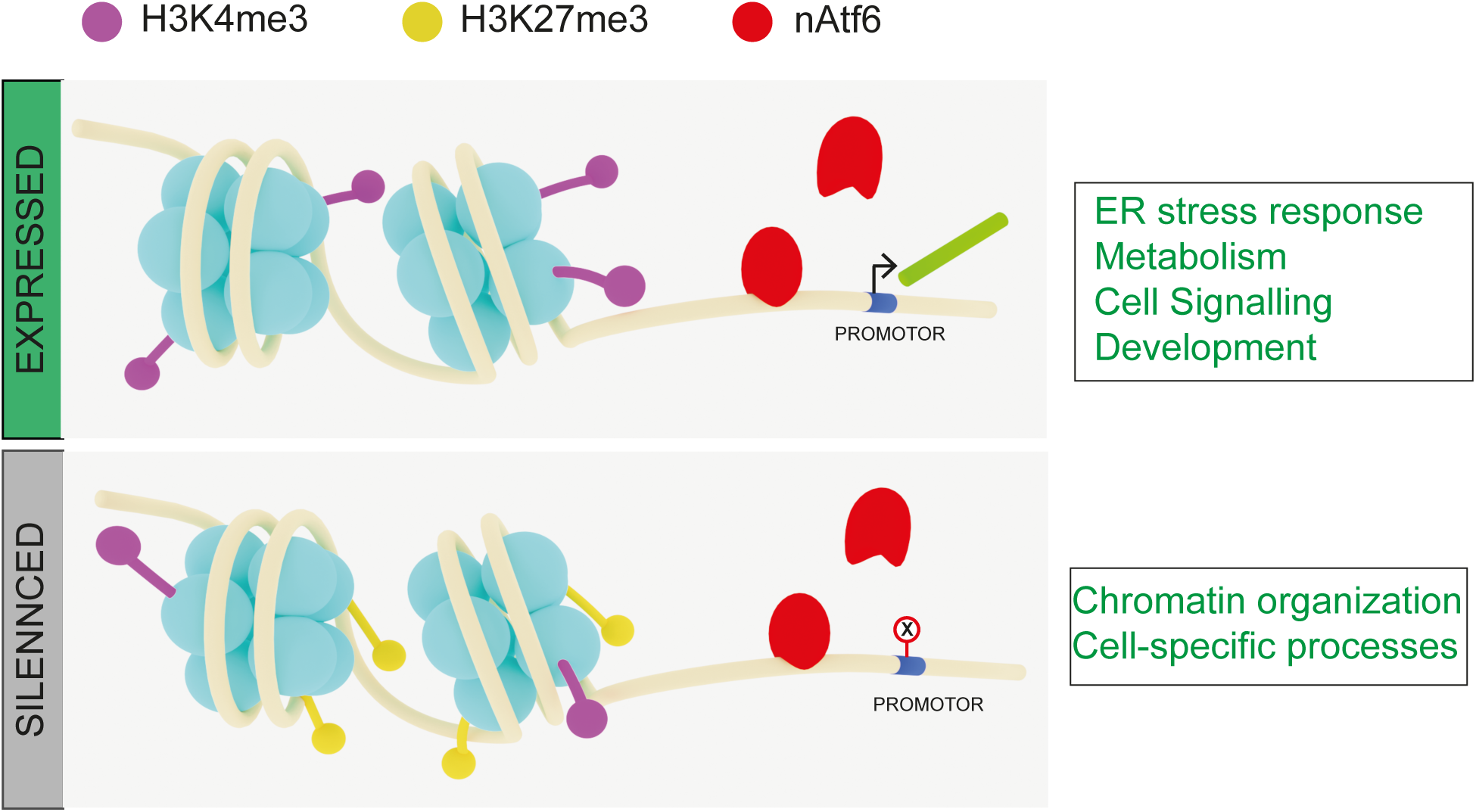
Epigenetic environment influences Atf6 mediated transcriptional output. Presence of nAtf6 by overexpression or toxicant exposure leads to differential expression of genes that fall within open chromatin with active transcription marks of H3K4me3 (purple) and functioning in proteostasis, metabolism and development. The CPATs can be expressed or repressed when nAtf6 (red) is present depending on the cellular environment and the presence or lack of co-factors (black). Contrastingly, toxicant and physiological stress could also change the epigenetic environment that could silence CPATs even in the presence of nAtf6 and being within open chromatin. This could be in part due to accumulation of repressive marks such H3K27me3 (yellow) that prevent the recruitment of the transcription machinery. These genes mainly function in chromatin organization or cell-specific processes that don’t express in hepatocytes. DNA in beige, histones in Cyan, newly generated mRNA lime green and TSS in blue.

These transcriptomic signatures in different cellular environments implicate co-factors or epigenetic modifications that direct Atf6 activity. The data presented here shows the CPATs that changed during mouse liver regeneration had a permissive epigenetic state in the quiescent liver. We speculate the absence of H3K27me3 allowed Atf6 to regulate CPAT expression, suggesting that accessible, active chromatin environment is conducive to Atf6 function as a transcriptional regulator (Figure 7). Many studies have shown that environmental exposures alter the epigenetic landscape [76, 77]; for instance, both arsenic and ethanol have been shown to alter epigenetic marks which are associated with mediating a toxicant dependent gene expression [78–80]. iAs has been suggested to change the methylation of histones and DNA [81, 82], and this could be a mechanism by which a distinct set of genes is differentially expressed in response to this toxicant. Further investigation of the epigenetic landscape and chromatin states in cells exposed to ER stressors will be important to identify such changes. An alternative model proposed by an interesting study of protein folding stress showed that sphingolipids activate distinct sets of Atf6 targets [15], suggesting that the mechanism by which Atf6 is activated dictates which targets are engaged. It would be of interest to determine if these different activation pathways also impact the chromatin state or epigenetic modifications on CPATs.

Toxicant exposure is predicted to have a very different transcriptional response than the stress-independent conditions for Atf6 activation; this is supported by our data showing that the transcriptomic response between iAs or EtOH exposure and genetic overexpression of Atf6 is substantially different. This could be explained by the involvement of other UPR pathways in response to toxicant exposure as well as the impact of other stress response pathways. In contrast, the early time point we analyzed in stress-independent conditions of Atf6 activation is the most reliable model to identify those genes that are targeted directly by Atf6. Interestingly, in the 78 hpf dataset where Atf6 was overexpressed, most of the DEGs are upregulated, and all but one CPAT genes (*bcat1*) are upregulated. This confirms that Atf6 primarily acts as a transcriptional activator; however, the subset of CPATs downregulated at later time points in the liver of *Tg(fabp10a:nAtf6-mCherry)* larvae, suggesting either that the downregulation of Atf6 targets require a prolonged engagement with Atf6 or a change in co-factors or chromatin structure that allows for gene repression. A recent study modeling the role of PERK in repressing transcription of Xbp1-Atf6 targets during prolonged stress exposure [83] suggests that crosstalk between pathways can shape the transcriptional response over time. Taken together, we conclude the UPR is a complex and interconnected network that depends on the stress intensity and cellular environment.

Previous studies have focused on identifying Atf6 targets using functional approaches in either Atf6 loss or overexpression models [29] [9] [43]. However, in most cases, the studies examine how such cells respond to stressed conditions using ER stress agents that severely impair major functions of the ER (i.e. tunicamycin or DTT). While these methods identify the UPR-related targets, they were not designed to identify Atf6 functional targets that participate in other biological processes. One study that explored other possible Atf6 functions using Atf6 knockout mesenchymal stem cells identified two sets of genes that are differentially expressed in the absence of Atf6 [84]: 145 ‘constitutive Atf6 responsive genes’ were downregulated by loss of Atf6 in standard culture conditions while another subset of 112 ‘induced Atf6 responsive genes’ were identified as those that failed to change their expression in response to tunicamycin treatment. The overlap between these two gene sets was only 6 genes, indicating that Atf6 regulates one set of genes in physiological conditions and another set – largely those targets involved in proteostasis - in response to ER stress. We compared these lists to our datasets; we found an overlap of only 1 gene between the constitutive targets and our putative Atf6 target genes. We speculate that different cell types and differences between *in vivo* approaches can significantly change the outcome of such studies.

Key caveats to our study to be considered are that the strict criteria used to identify putative Atf6 targets is based on previously described Atf6 binding sites and evolutionary conservation. Both of these can introduce error, in that the binding site predictions have not been thoroughly verified *in vivo* and the stringency of our evolutionary conservation criteria could exclude real Atf6 targets. However, given the lack of suitable antibodies to identify Atf6 genome occupancy through ChIPseq approaches, a logic based computational approach is an alternative. We are cognizant that the computational strategy used here could exclude some genes that may be relevant. This could be improved by including additional species, bridging the nodes on the evolutionary tree and altering the selection criteria from an exact match for the binding motifs to one that allowed substitution or spacing in the sequence. However, we reasoned that leniency could introduce false positives, and our preference was to increase the confidence that the CPATs are *bona fide* Atf6 targets. Generating reagents for genomic approaches to profile Atf6 binding sites and establishing how Atf6 target gene binding changes in response to environmental or physiological challenges is an important goal for the field.

A remarkable feature of the liver is its ability to regenerate after injury or resection. Many studies focus on the regulation of the cell cycle or metabolic processes that promote the regenerative response of the liver to partial hepatectomy (PH) [85, 86]. Here, we identified the novel involvement of Atf6 and CPATs during liver regeneration after PH in wild type mouse livers. These findings present a unique approach to identifying how Atf6 – and possibly other genes which have been exclusively only studied in the contexts of ER stress – may regulate physiological processes such as development and regeneration. Moreover, by using this novel model to study a UPR regulator, we uncovered an epigenetic pattern in the quiescent liver that may contribute to the ability of Atf6 to regulate target genes. The genes that are amenable for regulation after PH are devoid of the repressive mark, H3K27me3, are in open chromatin and are expressed at higher levels than CPATs that are not differentially expressed during regeneration. Interestingly, however, these silenced CPATs are also marked with H3K4me3 and in open chromatin, suggesting that they may be amenable to expression if the liver is exposed to other stimuli.

These findings have implications beyond the response to ER stress. It has long been known that the liver responds to a highly diverse range of environmental and physiological challenges; our findings suggest that, in part, this nuanced response is encoded in the epigenome. Moreover, these data indicate a broader range of processes that could be regulated by Atf6 and opens new areas of investigation for understanding what genes are regulated by this transcription factor and points to epigenetic changes as a mechanism contributing to target gene selection.

## ACCESSION NUMBERS

The datasets produced/used in this study have been deposited at Gene Expression Omnibus under the following accession numbers:

- **RNA-seq Data**:

○ **GEO Accession Number GSE130800** https://www.ncbi.nlm.nih.gov/geo/query/acc.cgi?acc=GSE130800
○ **GEO Accession Number GSE151291** https://www.ncbi.nlm.nih.gov/geo/query/acc.cgi?acc=GSE151291
○ **GEO Accession Number GSE125007** https://www.ncbi.nlm.nih.gov/geo/query/acc.cgi?acc=GSE125007
○ **GEO Accession Number GSE156419** https://www.ncbi.nlm.nih.gov/geo/query/acc.cgi?acc=GSE156419
- **ATAC-seq & ChIP-seq Data**

○ **GEO Accession Number GSE125006** https://www.ncbi.nlm.nih.gov/geo/query/acc.cgi?acc=GSE125006

## AUTHOR CONTRIBUTIONS

ARN and KCS conceived the idea and wrote the manuscript. ARN, PL and CZ carried out all the bioinformatic data analysis. ARN carried out all qPCR experiments and FM performed the Western Blot. KCS, ARN, PL, CZ and FM prepared the figures and analyzed data.

## ACKNOWLEDGEMENT

We gratefully acknowledge collaborative work with the NYUAD Genomics Facility and Bioinformatics Core, Tom Rutkowski for microarray data, Jaideep T.J. for the scientific illustration and to other members of the Sadler lab for sample preparation, fish maintenance and for their critical discussions.

## SUPPLEMENTARY METHODS

### Atf6 Target Gene comparisons

A list of predicted Atf6 target genes in humans was obtained from the R package ‘tftargets,’ from the Neph2012, TRED, TRRUST and Marbach2016 databases; this list of ‘Database Atf6 Targets’ was excluded from the final list of Atf6 target genes. Following this, we used Harmonize to identify the 116 genes curated by functional studies using Transfac (http://amp.pharm.mssm.edu/Harmonizome/gene_set/ATF6/TRANSFAC+Curated+Transcription+Factor+Targets). Due to minimal overlap with this list, as well as lack of canonical Atf6 target genes this list was excluded as well.

### Western Blot

Zebrafish 5 dpf embryos of *Tg(fabp10a:nls-mCherry)* [67] *from* WT siblings and *Tg(fabp10a:nAtf6-mCherry;cmlc2-GFP)* were dissected using syringes needle according to our previously published protocol. Protein extraction was performed on 20 livers/condition in 50ul Lysis buffer, and sonicated with 2 second pulse for 30 times (amplitude 30%). The proteins were clarified in refrigerated centrifuge at 4° C for 15 min with speed >10.000g. Quantification with QuBit protein Kit was performed, and 15ug/lane of proteins was loaded on denaturating 10% gel (BIORAD cast fast). Then standard run (80-150V) and transfer (400 mA for 1:30 hours). Blocking Milk 5% in TBS-T for 1h and Primary Abs O.N. 4 C° (Mouse, α-mCherry 1:1000 in Milk 5%, and Rabbit α-H3 1:1000 in Milk 5%, Santa Cruz Biotechnology, 10809). Washing 3 times in TBS-T and secondary 1:2000 1-2h was performed subsequently. Development was performed in BioRad ChemiDoc with ECL Thermo-Pierce (Low sensitivity reagent), or ECL Clarity BioRad (Mid sensitivity reagent), or ECL Select Amersham-GE HealtCare (High sensitivity reagent).

## SUPPLEMENTARY FIGURE LEGENDS

**Supplementary Figure 1.**
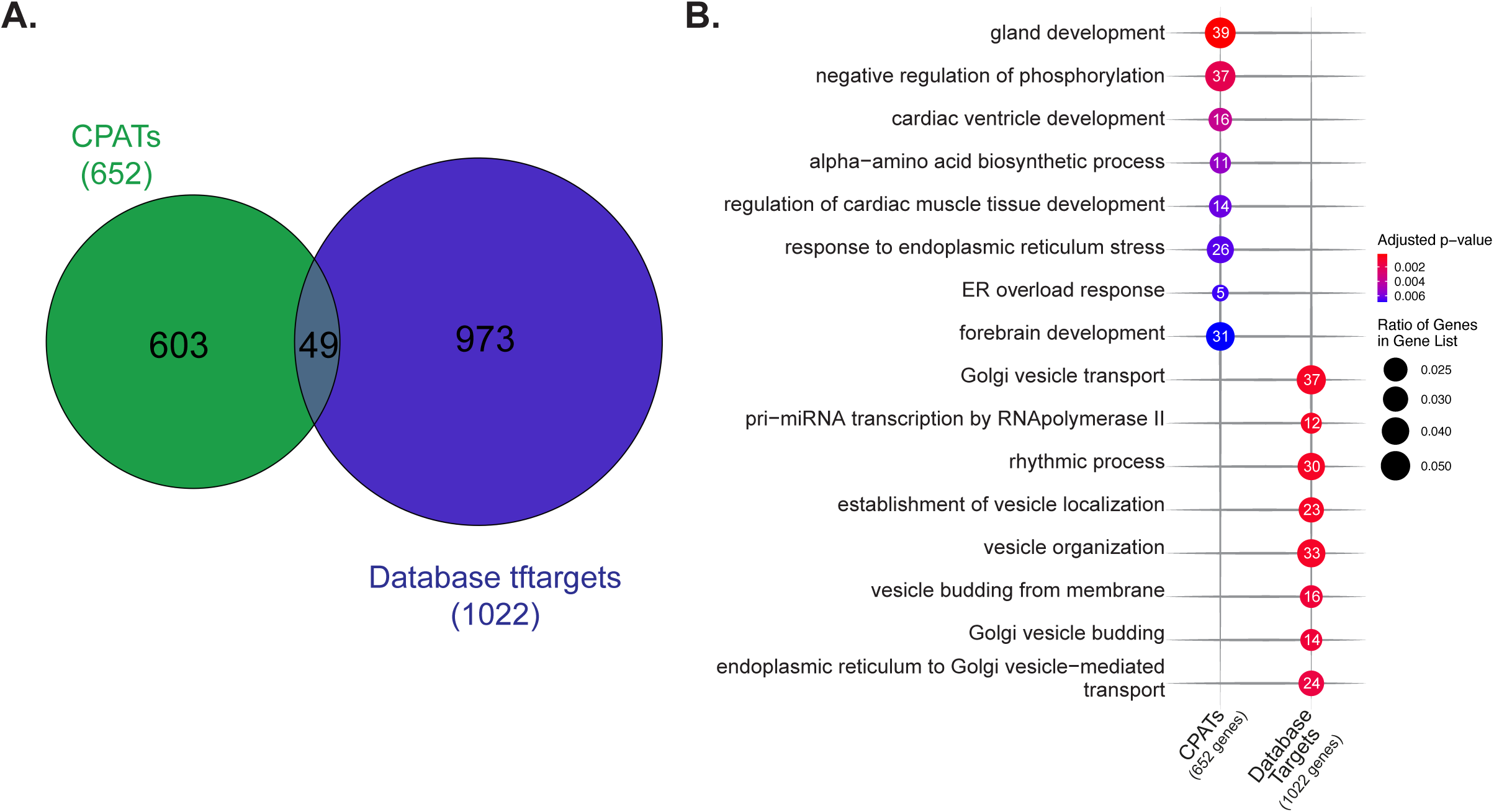
Atf6 putative target has minimal overlap with database targets. (A) Venn diagram depicting the overlap between database targets acquired through tftargets (purple) and CPATs (green). (B) Gene ontology-based biological process overrepresentation analysis of database targets and CPATs. Statistically significant results are plotted with point size depicting fraction of genes in each gene list annotated to each GO term, with point numbers showing the exact number of genes annotated.

**Supplementary Figure 2.**
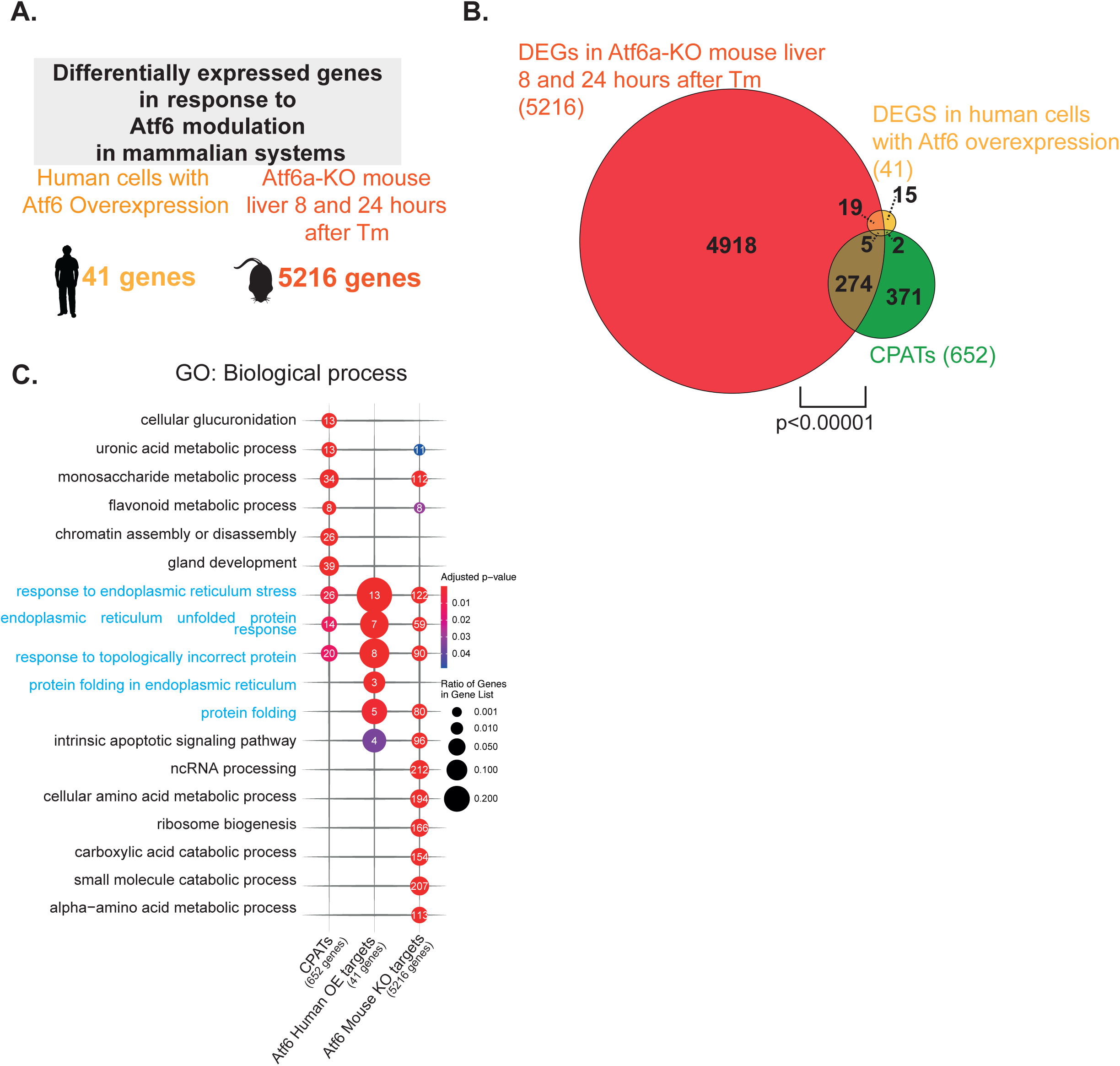
Modest representation of CPATs in Human HEK293 cells overexpressing nAtf6 and in livers from Mouse Atf6a knockout exposed to Tunicamycin. (A) Schematic showing the number of genes differentially expressed in each publicly available dataset. (B) Venn diagram showing overlap of CPATs (green) with DEGs in Human HEK293 cells overexpressing nAtf6 (yellow) and in livers from Mouse Atf6a knockout exposed to Tunicamycin (red).p<0.0001 by Chi-square test. (C) Gene Ontology based biological process enrichment analysis of CPATs and DEGs from microarray data of nAtf6 overexpression Human cells and Atf6 knockout mice.

**Supplementary Figure 3.**
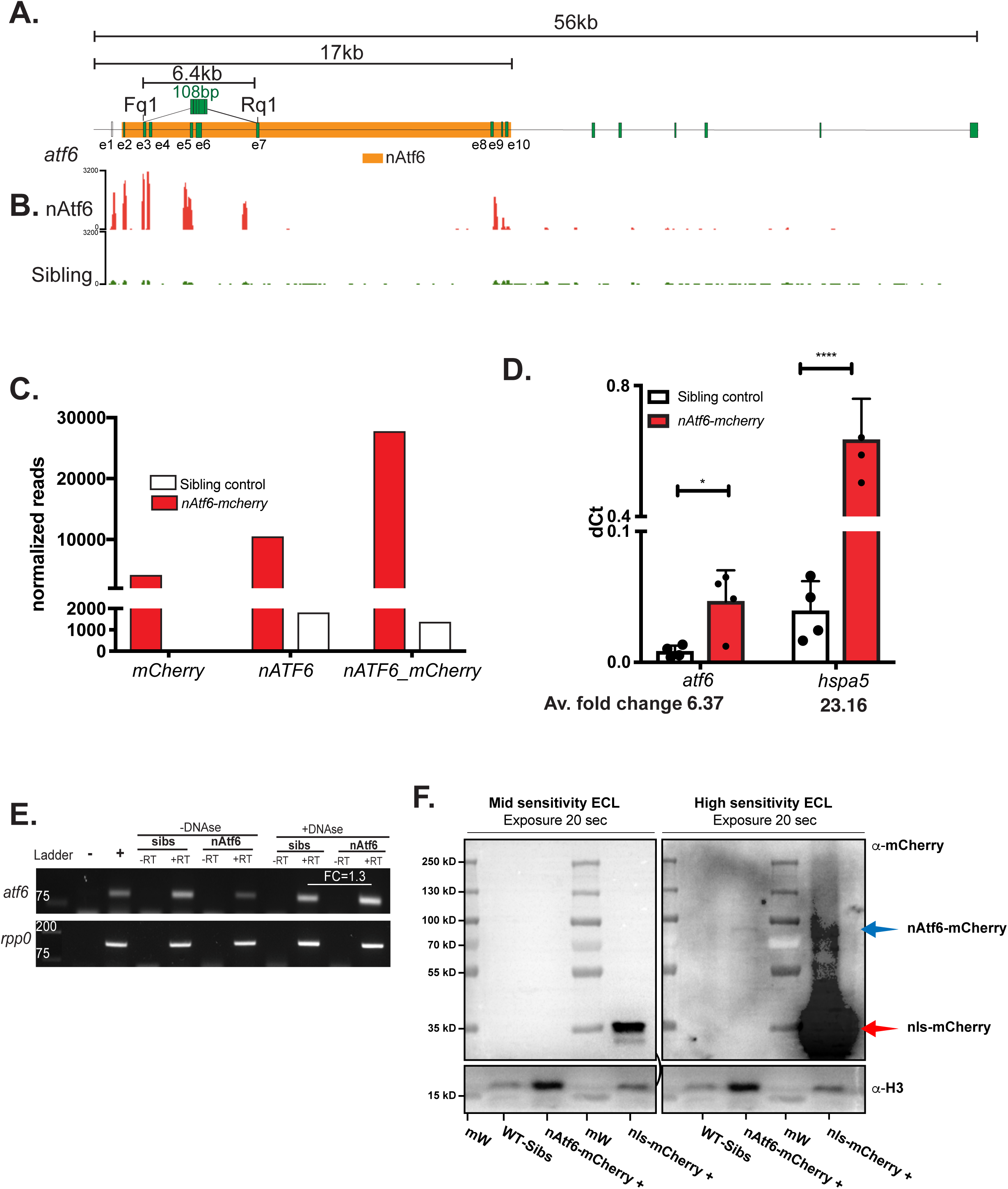
nAtf6 overexpression in the liver of *Tg(fabp10a:nAtf6-mcherry;cmlc2:GFP)-C* zebrafish. (A) Representation of *atf6* genomic region. (B) Browser view of reads detected by RNAseq from nAtf6 overexpression (red) and siblings (green) across gene region of Atf6. (C) Bar plot showing normalized reads of cherry, nAtf6 and nAtf6-mcherry junction detected in controls and *Tg(fabp10a:nAtf6-mcherry;cmlc2:GFP)-C* livers. (D) qPCR analysis of RNA isolated from the liver of 5 clutches of zebrafish larvae from *Tg(fabp10a:nAtf6-mcherry;cmlc2:GFP)-C* and non-transgenic siblings on 5 dpf. *atf6* and *hspa5* (*bip*) were normalized to *rpp0* and the dCt was plotted; with fold change in transgenics normalized to siblings indicated. * p-value <0.05 and **** p-value <0.00005 by Students t-test. (E) Agarose Gel showing Atf6 mRNA detected in both sibling and nAtf6-mcherry in samples pretreated with DNAse and without as well as with reverse transcriptase and without to confirm that there was no DNA contamination. *rpp0* was used for loading normalization. (F) Top-left panel shows α-mCherry developed with mid sensitivity reagent and exposed for 20 sec. Top-right panel shows α-mCherry developed with high sensitivity reagent and exposed for 20 sec. The difference of detectability for nAtf6-mCherry and nls-mCherry proteins are highlighted by arrows: blue arrow indicates the nAtf6-mCherry fusion protein around 90 kD. The expected fusion protein is around 70kD, and is expected to shift for multiple glycosylation of the nuclear fraction of Atf6 as well-reported in literature; red arrow indicated nls-mCherry protein (∼35 kD) that is detectable with mid sensitivity ECL reagent on top-left panel and becomes overexposed in top-right panel with high sensitivity ECL reagent. Bottom panels show α-H3 used for loading normalization and developed with low sensitivity reagent and exposed for 15 seconds.

**Supplementary Figure 4.**
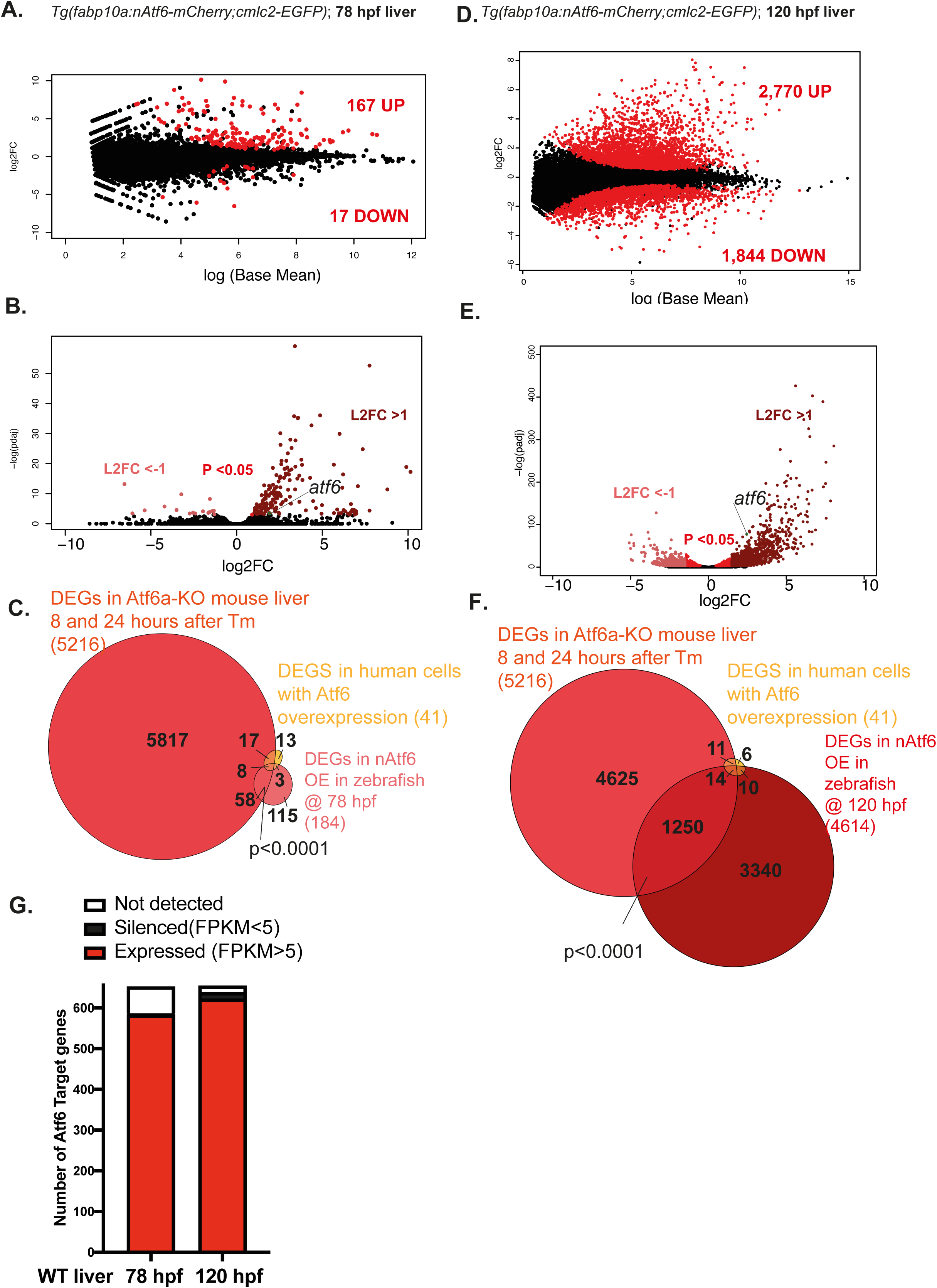
nAtf6 overexpression in zebrafish livers shows both transcriptional activation and repression of genes. MA plot of RNAseq from *Tg(fabp10a:nAtf6-mcherry;cmlc2:GFP)-C* livers relevant to non-transgenic siblings at 78 hpf (A) and 120 hpf (D). Red dots indicate genes that have padj<0.05 and are considered significant. Volcano plot showing significantly upregulated genes (maroon) and downregulated genes (pink) by nAtf6 overexpression at 78 hpf (B) and 120 hpf (E). Red dots indicate ge\nes that are significant but the fold change is minimal. Venn diagram showing overlap of DEGs from microarray data of nAtf6 overexpression Human cells and Atf6 knockout mice with RNAseq from nAtf6 overexpression at 78 hpf (C) and 120 hpf (F). (G) Bar plot depicting the number of CPATs expressed in the liver of non-transgenic siblings at 78 hpf and 120 hpf based on a cut off of FPKM>5 as expressed, FPKM<5 as silenced; some genes were not detected in any dataset.

**Supplementary Figure 5.**
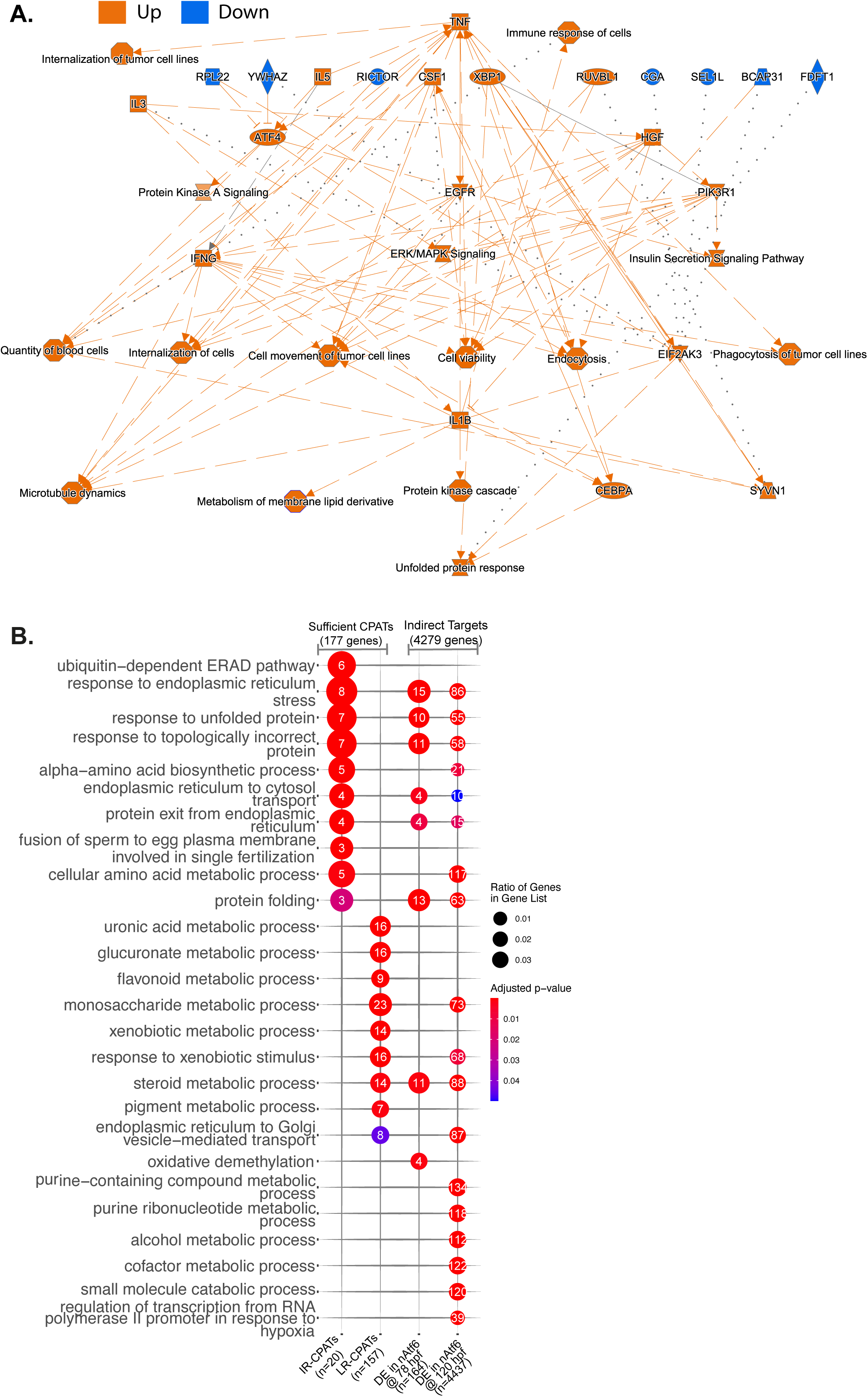
nAtf6 overexpression leads to activation of other transcription factors that are involved in functions other than Proteostasis. (A) Summary Plot of pathways from IPA affected due to nAtf6 overexpression in livers at 120hpf. (B) Gene Ontology based biological process enrichment analysis of sufficient CPATs and Indirect Atf6 targets.

**Supplementary Figure 6.**
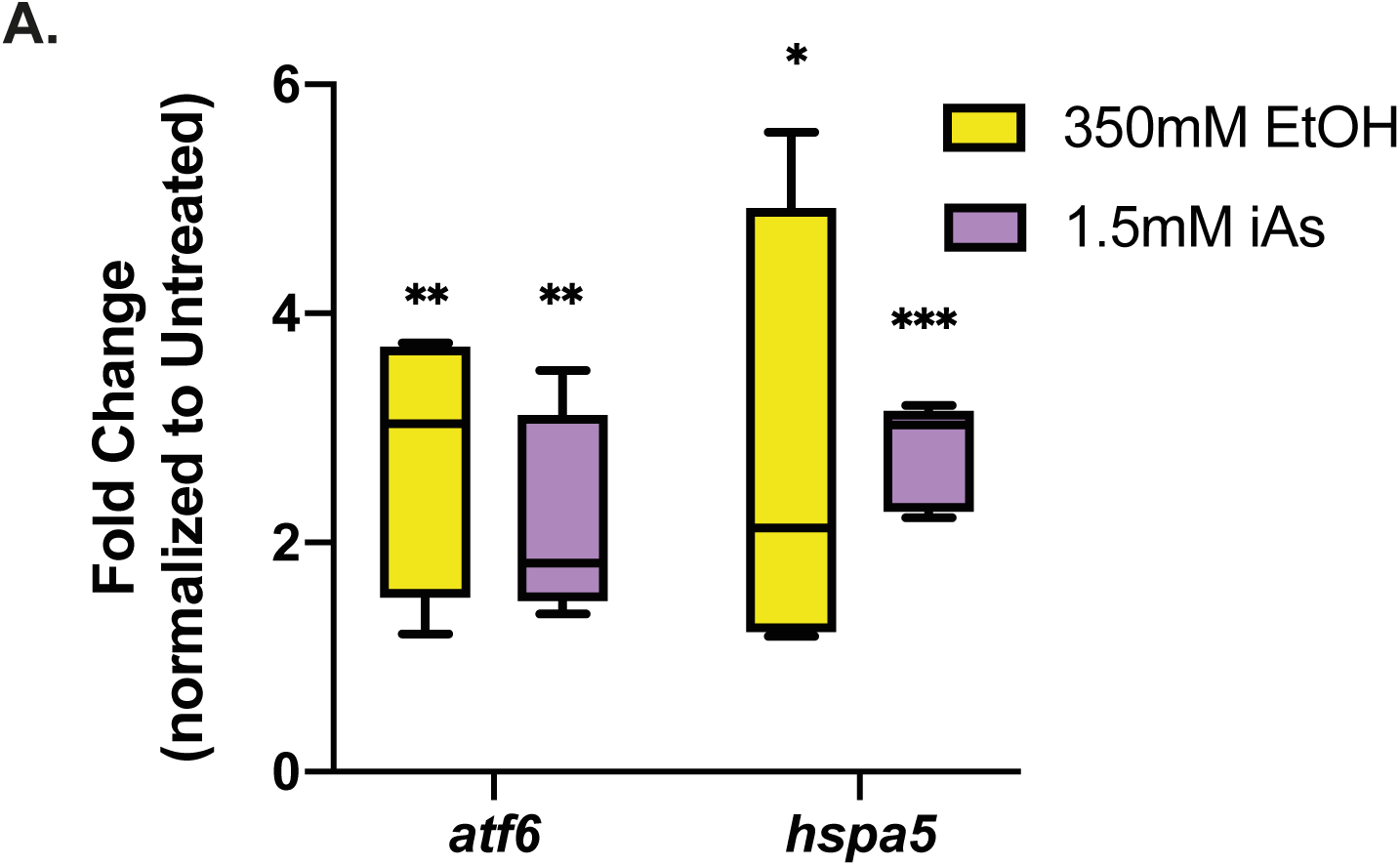
iAs and EtOH treatment leads to upregulation of *atf6* and *hspa5*. Expression Analysis by RNA-seq on iAs-treated and EtOH-treated zebrafish livers at 120 hpf showing foldchange of *atf6* and *hspa5* in comparison to untreated and normalized to housekeeping gene *rpp0*. ***p-value<0.0005, **p-value<0.005, *p-value<0.05 by Students t-test.

**Supplementary Figure 6.**
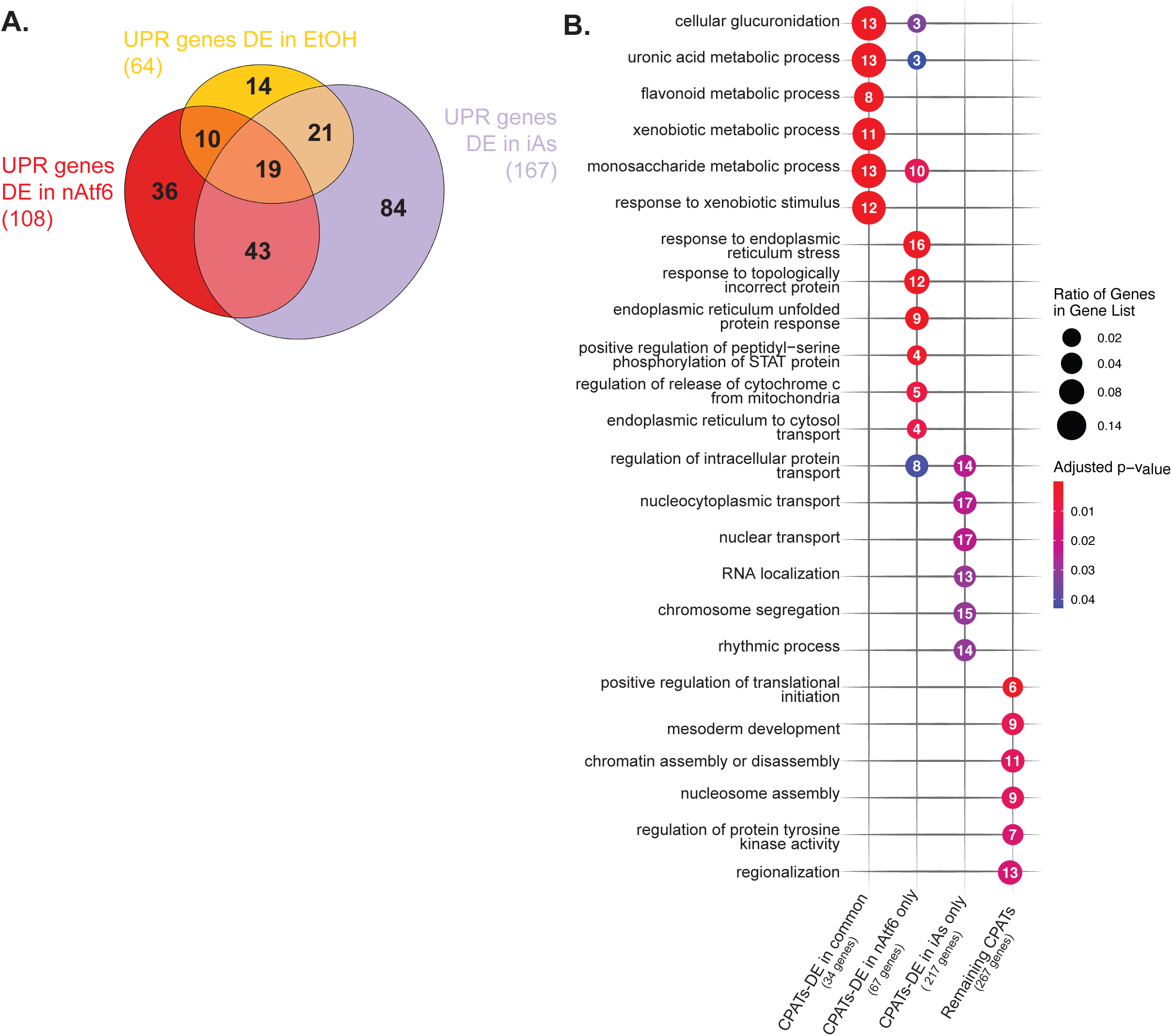
CPATs and UPR genes show stress dependent distinct patterns of expression. (A) Venn diagram depicting the distribution of UPR genes differentially expressed in all conditions: nAtf6, EtOH and iAs. (B) Gene Ontology based biological process enrichment analysis of CPATs DE in common and unique in nAtf6 and iAs.

**Supplementary Figure 7.**
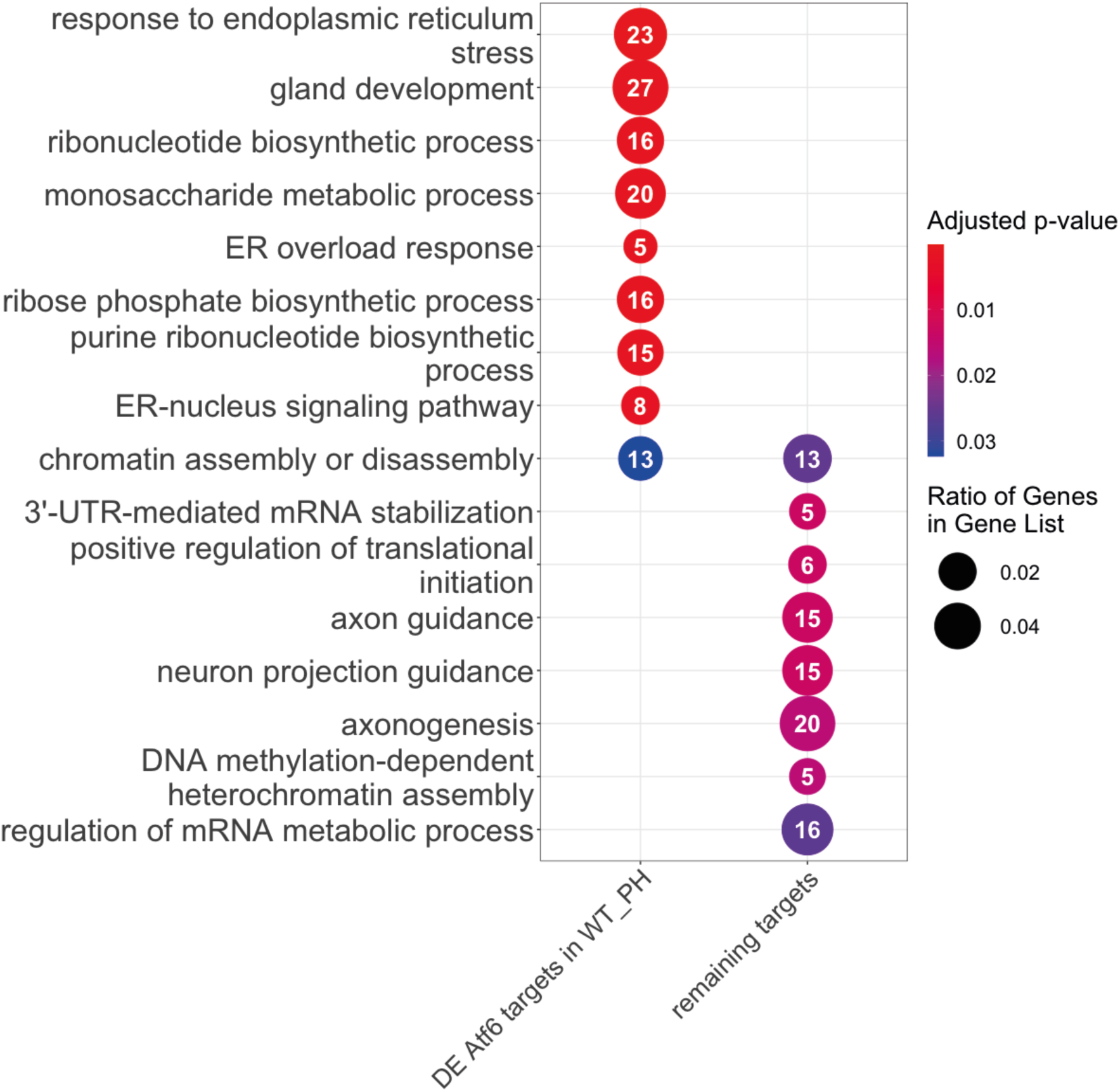
DE-CPATs during regeneration are involved in proteostasis and metabolic processes while those that do not change function in chromatin organization and cell-specific functions. Gene Ontology based biological process enrichment analysis of CPATs that are DE and those that do not change in regenerating mouse liver.

## SUPPLEMENTARY TABLES

**Supplementary Table S1.** List of qPCR primers used in the study.

**Supplementary Table S2.** Summary of all Zebrafish Ensembl IDs and Gene names of Atf6 gene lists used for the study: NFY consensus sites (CCAAT/ATTGG), Atf6 binding site(CCACG) and Atf6 RE (TGACTG)

**Supplementary Table S3.** Summary of CPATS comprising of Zebrafish Ensembl IDs and Gene names and corresponding Ensembl IDs of homologs in Human, Mouse, Zebrafish

**Supplementary Table S4.** Summary of all Zebrafish Ensembl IDs and Gene names of database and publicly available datasets used to validate the CPATs: tftargets, genes differentially expressed in nATF6 over expression in human HEK293 cells, in ATF6 knockout mice exposed to Tunicamycin and the overlapping genes between them.

**Supplementary Table S5.** Summary of UPR GO and Reactome dataset used for the study: Reactome UPR, GO 0034976 ER Stress and GO 0006986 Unfolded Protein.

**Supplementary Table S6.** Table of putative Atf6 target genes and UPR genes that are differentially expressed or not expressed across all three RNAseq datasets (nAtf6, EtOH and iAs).

**Supplementary Table S7.** Table of all four RNA seq datasets (nAtf6-78, nAtf6-120, EtOH and iAs) consisting of Gene name, Zebrafish Ensembl ID, log2Foldchange, adjusted p-value and whether they are Atf6 target/ UPR genes and if differentially expressed in other corresponding datasets and the presence of NFY, Atf6 binding site or Atf6 RE in zebrafish.

**Supplementary Table S8.** Summary of gene expression analysis by qPCR on CPATs generated from multiple clutches pretreated with either DMSO or Ceapins and then co-exposed to iAs.

**Supplementary Table S9.** Summary of DEGS during wild type mouse liver regeneration consisting of Mouse Ensembl IDs, gene names, whether they are CPATs and at which timepoint they are differentially expressed.

**Supplementary Table S10.** Summary of DE-CPATs during wild type mouse liver regeneration divided into clusters based on their expression.

